# Genetic variation associated with plastic and homeostatic growth responses to drought in Arabidopsis

**DOI:** 10.1101/2021.05.07.443185

**Authors:** Ángel Ferrero-Serrano, Sarah M. Assmann

## Abstract

Natural genetic variation influences plant responses to environmental stressors. However, the extent to which such variation underlies plastic versus homeostatic response phenotypes deserves more attention. We quantified the extent of drought-induced changes in leaf area in a set of Iberian Arabidopsis accessions and then performed association studies correlating plasticity and homeostasis in this phenotype with genomic and transcriptomic variation. Drought-induced plastic reductions in relative leaf area typified accessions originating from productive environments. In contrast, homeostasis in relative leaf area typified accessions originating from unproductive environments. Genome-Wide Association Studies (GWAS), Transcriptome Wide Association Studies (TWAS), and expression GWAS (eGWAS) highlighted the importance of auxin-related processes in conferring leaf area plasticity. Homeostatic responses in relative leaf area were associated with a diverse gene set and positively associated with a higher intrinsic water use efficiency (WUE_i_), as confirmed in a TWAS metanalysis of previously published δ^13^C measurements. Thus, we have identified not only candidate “plasticity genes” but also candidate “homeostasis genes” controlling leaf area. Our results exemplify the value of a combined GWAS, TWAS, and eGWAS approach to identify mechanisms underlying phenotypic responses to stress.

**Highlight:** Information on phenotype, genotype, and transcript abundance is integrated to identify candidate plasticity and homeostasis genes and processes associated with local adaptation to drought stress in Arabidopsis accessions of the Iberian Peninsula.

## Introduction

In the Italian novel The Leopard, one of the main characters famously conveys the idea that ‘Everything must change so that everything can stay the same.’ Often, there is a cost associated with change that resonates with the plastic developmental responses observed in plants when faced with stress (Bradshaw, 1965; West-Eberhard, 2003). Plasticity marks a departure from a homeostatic state in response to an environmental cue, often followed by “allostatic” responses to return to a homeostatic state (Fig. 1). These allostatic responses involve a myriad of physiological and developmental regulatory processes that stabilize critical aspects of plant physiology. Under repeated stress there is an associated metabolic and physiological cost associated with departure from the optimal state that is not completely alleviated upon return to homeostasis. These retained detrimental effects are known as allostatic load (McEwen and Stellar, 1993), with important implications for fitness and survival from an evolutionary standpoint.

**Fig. 1.**
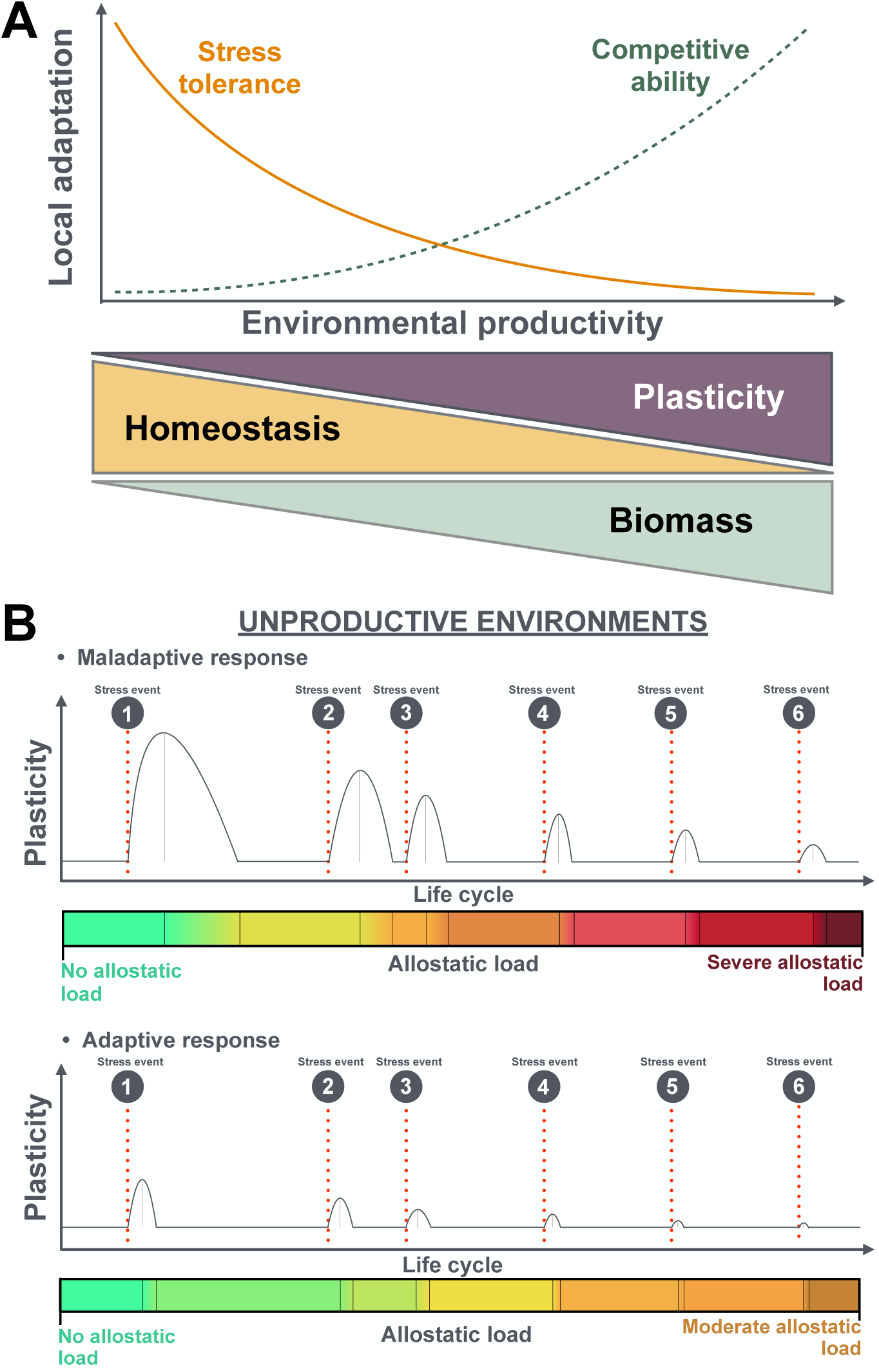
Conceptual framework for the adaptive role of plastic and homeostatic responses across productivity gradients in the local environment. (A) Schematic illustration representing the balance *sensu* Grime (1974, 1977), between stress tolerance in lower productivity, stress-prone environments, and competitive ability in and homeostasis in plants. In less productive environments, homeostatic responses to stress and lower relative growth rates result in lower biomass production, which is tuned in to the lower resources that are typically available in their environment and favors adaptation to those environments. In contrast, in productive environments, in which competition becomes more intense, plastic responses are favored from an adaptive standpoint. (B). In unproductive environments, frequent stress events can increase the allostatic load in individuals adapted to high-productivity environments where plastic responses are beneficial. During the lifetime of an individual, this results in a maladaptive response that increases allostatic load. In contrast, individuals adapted to unproductive environments exhibit less plastic (more homeostatic) responses to repeated stress that reduces their allostatic load over their lifetime.

Stress often elicits plastic responses in plant developmental traits (Dorn *et al*., 2000; van Kleunen and Fischer, 2005; Forsman, 2015). However, simply because a given trait is plastic does not mean that this plasticity is adaptive (Ghalambor *et al*., 2007). Plants that are not adapted to stress may exhibit plastic responses that are maladaptive. Under such conditions, homeostasis or lack of plasticity may, in turn, be adaptive (Fig. 1; Ferrero-Serrano and Chakravorty, 2023). This study aims to explore the genetic basis of both plasticity and homeostasis imposed by recurrent drought events during the vegetative development of *Arabidopsis thaliana*.

Drought is among the strongest factors driving adaptation in plants (Siepielski *et al*., 2017; Exposito-Alonso *et al*., 2019). Since climate projections predict drought events to become more frequent, intense, and unpredictable (Donat *et al*., 2016), we can expect that drought episodes will become increasingly more difficult for plants to adapt to. For this reason, understanding the genetic basis and adaptation potential of phenotypic traits and their plastic or homeostatic responses to drought is paramount for ameliorating climate change impacts in natural and managed ecosystems.

The Iberian Peninsula is a drought-prone region particularly vulnerable to the effects of climate change on precipitation (Vicente-Serrano, 2006a; Vicente-Serrano, 2006b; Coll *et al*., 2016; Páscoa *et al*., 2017). The Iberian Arabidopsis population is particularly valuable because its accessions originate from a broad range of environments (Lobo *et al*., 2001; Rey Benayas and Scheiner, 2002), and previous studies of this population have provided considerable insight into Arabidopsis evolution and development (Picó *et al*., 2008; Montesinos *et al*., 2009; Gomaa *et al*., 2011; Méndez-Vigo *et al*., 2011; Montesinos-Navarro *et al*., 2011; Picó, 2012; Manzano-Piedras *et al*., 2014; Wolfe and Tonsor, 2014; Vidigal *et al*., 2016; Exposito-Alonso *et al*., 2018a; Marcer *et al*., 2018; Castilla *et al*. 2020). In the present study, we utilize GWAS, Transcriptome-Wide Association Studies (TWAS), and Expression Genome-Wide Association Studies (eGWAS) to identify natural genetic and transcript abundance variation underlying both plastic and homeostatic leaf area responses to drought in Iberian Arabidopsis accessions. Leaf area is a vital trait in relation to growth: leaf area production determines the total amount of light intercepted by plants, carbon assimilation, and the extent of evaporative water loss. We uncover an adaptive homeostatic growth strategy in accessions from less productive environments, address its genetic basis, and conclude that phenotypic homeostasis is under genetic control and of evolutionary importance, just as previously argued for plasticity (Bradshaw, 1965; Via *et al*., 1995).

## Materials and Methods

### Plant materials and phenotyping

Seeds for 43 Iberian accessions (Table P1 for the list of accessions) were cold-stratified for four days at 4°C to synchronize germination. Plants were grown in 10 cm square pots containing Metro-mix 360 potting mixture (Sun Gro Horticulture Canada Ltd) in a greenhouse with an average daytime 500 μmol m^−2^s^−1^ photosynthetic photon flux density (PPFD) (8h light: 16h dark, with light supplied as natural daylight supplemented with 1000W metal halide lamps) and average day/night temperatures of 22°C /20°C.

For each Iberian accession, we grew one plant per pot, with five replicates for each of the control and drought treatments (10 plants for each individual accession), for a total of 430 plants (43 accessions × (five control + five droughted plants)). This is a greater number of replicates than for many association studies (Seren *et al*., 2020). Both control and drought treatment were well-watered for the first 28 days, maintaining the soil relative water content at field capacity as measured by monitoring pot weight daily and adding water when necessary. After 28 days, soil in the drought treatment was allowed to dry down. These pots were then rewatered to soil saturation once every 10 days for a total of 14 drought cycles until the conclusion of the experiment, 175 days after seeds were sown. Here, as in the natural environments we sought to mimic, the soil water content was not adjusted based on the size of individual plants. Our protocol simulated the natural recurring cycle of rainfall and drought. Such cycles are often used in studies of natural variation (i.e. Exposito-Alonso *et al*., 2018b; Exposito-Alonso *et al*., 2019) and in field trials on crop cultivars (IRRI, 2021). In contrast, the well-watered treatment was maintained near field capacity throughout the experiment. Plants were spatially randomized within the greenhouse every week throughout the experiment. At the end of the experiment, nine accessions within the drought treatment with mortality rates higher than 40% (i.e., at least two of the five replicates; Table P1) were excluded from downstream analyses.

Photographs of the rosettes were taken at the time of flowering and subsequently analyzed using ImageJ (http://imagej.nih.gov/ij/) to determine the projected leaf area for each accession under both well-watered and drought treatments (Fig. S1; Table P1).

### Determination of relative leaf area and plasticity

Relative leaf area (rLA) is defined here as the leaf area (LA) at the time of flowering for each accession under each experimental condition (either well-watered or drought) relative to all the accessions under the same condition. The relative leaf area of a given accession under well-watered conditions (rLA_i (well-watered)_) is thus calculated as:

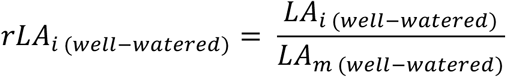

where LA_i (well-watered)_ is the leaf area of any individual accession, and LA_m (well-watered)_ is the average leaf area for all accessions grown under the same conditions. Similarly, the relative leaf area under drought (rLA_i (drought)_) for each accession is calculated as:

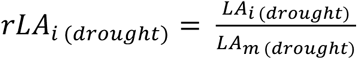

where LA_i (drought)_ is the leaf area of any individual accession under drought and LA_m (drought)_ is the average value for all accessions when grown under drought. A high rLA reflects greater vegetative growth of any given accession relative to the set of accessions utilized in this study. The use of rLA, rather than actual values, was essential to allow a normalized comparison of the leaf area of each accession between the watering regimes. Reaction norms based on rLA were employed to quantify plastic changes in relative leaf area between the two treatments (well-watered vs. drought). Leaf area plasticity indexes were calculated based on an established definition (Weijschedé *et al*., 2006). The leaf area plasticity index to drought (PI) was calculated for each accession as follows:

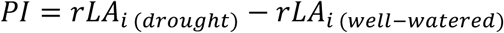

A positive leaf area plasticity index does not necessarily imply that the leaf area for that accession was higher under drought, but rather that under drought, a given accession displays a higher rLA relative to the rest of accessions in the study.

### Genome-wide association study

Genome-Wide Association Studies (GWAS) identify genetic variants associated with natural variation in a phenotypic trait of interest. We used a GWAS approach to identify polymorphisms associated with natural variation in rLA and its plasticity. The online tool GWAPP (http://gwas.gmi.oeaw.ac.at/) was employed using a linear regression model (Seren *et al*., 2012). The linear regression model does not address the confounding effects of population stratification, family structure, and cryptic relatedness (Price *et al*., 2010); however, these issues are minimized by our choice of a regional (Iberian) population (Frachon *et al*., 2018; Tabas-Madrid *et al*., 2018; Frachon *et al*., 2019). Nonetheless, we demonstrate that the candidate SNPs discussed here were also significant after the application of a mixed model approach implemented either using GWAPP (Seren *et al*., 2012) or GWAS-Flow (Freudenthal *et al*., 2019) (Figs S2, S3, and S4; Tables G1-G6), providing further confidence in our results.

We filtered out low-frequency variants (minor allele frequency (MAF) < 10%) to focus on variants less likely to be undergoing negative selection or to be spurious SNPs from sequencing errors. We obtained gene model annotation from the TAIR11 genome release. We defined the ‘consensus’ transcript variant as the variant producing the longest transcript. We predicted the effect of every SNP using SnpEff (Cingolani *et al*., 2012). We focused on SNPs within gene transcriptional units, including introns, untranslated regions at the 5′ and 3′ ends, and open reading frames, as defined in Araport11 (Cheng *et al*., 2017). We also included the promoter region, defined as 5 kb upstream of the most distal transcription start site. We imposed a stringent arbitrary significance threshold corresponding to a score of four (*P* < 0.0001; Fig. S5A-E).

Using the results from GWAS obtained from the association with rLA, we obtained reaction norms that reflect plastic or homeostatic patterns of genome-wide associations of these traits in response to drought. The association between a phenotype and any given SNP was designated as homeostatic if it was found significantly associated with that trait under both conditions (well-watered and drought).

### Transcriptome-wide association

Natural variation in gene expression is an important mechanism of local adaptation (Des Marais *et al*., 2012; Lasky *et al.,* 2014). Transcriptome-Wide Association Studies (TWAS) identify variation in transcript abundance associated with observed variation in phenotypic traits (Gusev *et al*., 2016; Kremling *et al*., 2019). TWAS analysis does not suffer from the limitations imposed by false non-causal synthetic associations due to linkage disequilibrium (Li *et al*., 2021) that are typical of GWAS (Korte and Farlow 2013). Also, TWAS evaluates continuous variation in transcript abundance rather than the discrete presence/absence of SNPs with (potentially) additive or synergistic effects on a continuous phenotype. We retrieved the transcriptome data from rosette leaves of 727 Arabidopsis accessions from the GEO dataset with accession number GSE80744 and SRA study SRP074107 (Kawakatsu *et al*., 2016) and subsetted the 35 Iberian accessions that overlapped with the 43 Iberian accessions used in our study. Data wrangling was conducted utilizing the dplyr (Wickham *et al*., 2015) and tidyr (Wickham and Henry, 2018) packages. Spearman’s rank correlation coefficients between the variation in individual phenotypic values (relative leaf area (rLA_i_), plasticity index (PI), and intrinsic water use efficiency (WUE_i_) from Dittberner *et al*. (2018; Tables P1 and P2)), and individual transcript abundance values within the study population, were calculated using the correlation function of the Hmisc package (Harrell Jr and Dupont, 2019). The stronger the association of the two variables, the closer the Pearson’s correlation coefficient, r_s_, will be to either +1 or −1. We imposed a threshold based on this correlation coefficient (|r_s_| ≥ 4.0). We considered this threshold to be stringent after inspection of the distribution of the results (Fig. S5F-J). We include the r_s_ and corresponding *P-*values in Tables T1 and T2.

Using the results from TWAS, we derived reaction norms to reflect homeostatic patterns of transcriptome-wide associations of rLA in response to drought. Any given transcript was designated homeostatic if found significantly associated with rLA (either positively or negatively; |r_s_| ≥ 4.0) under both conditions (well-watered and drought).

### eGWAS

Natural variation in transcript abundance can also be used to identify candidate SNPs (expression SNPs or eSNPs) that control transcript abundance in an Expression Genome-Wide Association Study (eGWAS). eGWAS was conducted by compiling the transcript abundance values of a given gene of interest, extracting the transcript abundance for that particular gene in the 727 Arabidopsis accessions from the GEO dataset with accession number GSE80744 and SRA study SRP074107 (Kawakatsu *et al*., 2016), then subsetting the 665 accessions included in the 1001 Genomes project (The 1001 Genomes Consortium, 2016). Then, the online tool GWAPP (http://gwas.gmi.oeaw.ac.at/) was employed using a linear regression model (Seren *et al*., 2012) to identify candidate regulatory variants. Regulatory variants are either cis- or trans-acting, depending on the location of the genetic variant. cis-eSNPs were defined as those located within the ORF, introns, UTRs, or the promoter region of the regulated gene. Trans-eSNPs were defined as those located within the ORF, introns, UTRs, or promoter region of a gene different from the gene transcriptionally regulated by that genetic variant.

### Analyses

AraCLIM (https://github.com/CLIMtools/) is a database that describes the local environment of the Arabidopsis accessions sequenced as part of the 1001 Genomes Project (Ferrero-Serrano and Assmann, 2019; Ferrero-Serrano et al., 2022). We obtained from AraCLIM the values for MOD17A2 (Net Primary Productivity in the spring season) (Heinsch *et al*., 2003; Running and Zhao, 2015) at the collection site of each accession and evaluated the correlation with the phenotypes obtained in our experiment.

WUE_i_ is defined as the ratio between instantaneous carbon gain and transpirational water loss, typically calculated per unit of leaf area (Condon *et al*., 2002). To integrate information from previously published studies on the Iberian Arabidopsis population, we retrieved available data on natural variation in intrinsic water use efficiency (WUE_i_, δ^13^C), previously recorded by Dittberner *et al*. (2018). During photosynthesis, plants discriminate against ^13^CO_2_ in the reaction center of *Ribulose 1, 5-biphosphate carboxylase/oxidase* (*RuBisCO*). As stomata close, minimizing water loss in response to drought, the discrimination against ^13^CO_2_ decreases because the internal CO_2_ concentration (C_i_) becomes unavoidably reduced. As a result of their reduced C_i_, plants with greater WUE_i_ experience lower CO_2_ concentrations at the sites of carboxylation (C_c_) and have less negative δ^13^C values.

All statistical analyses and figures were produced using R (Team, 2013). The code and data to reproduce the analysis, figures, and tables are available at https://github.com/AssmannLab and Zenodo. Gene Ontology (GO) biological process analysis (Tables GO1-GO10) was performed using the open-source software ShinyGO v0.61: http://bioinformatics.sdstate.edu/go60/ (Ge *et al*., 2020) with the following settings: the search species was “*Arabidopsis thaliana*,” the *P*-value cutoff (FDR) was 0.05, and the number of most significant terms to show was ten. To replicate these results, the lists of genes that were input into these analyses based on significance thresholds can be selected from Tables G1-G6, T1, and T2. Each analytical technique used in this study has assumptions and limitations, which we discuss in Document S1. The code and data necessary to reproduce all analyses and figures in this study are available on GitHub (https://github.com/AssmannLab).

## Results

### Natural variation in leaf area and plasticity in response to long-term cyclical drought

To quantify phenotypic variation in Iberian Arabidopsis accessions exposed to drought, we conducted a common garden experiment (Fig. 2A). We used image analysis to derive rosette leaf area under well-watered or drought conditions (Figs 2B, S1; Table P1). As expected, we observed a significant (*P* < 0.001) reduction in leaf area resulting from drought in most genotypes, as evidenced by the phenotypic reaction norms. In contrast, a small number of accessions exhibited constant or increased leaf area under drought (Fig. 2B; Table P1), demonstrating that a reduction in leaf area in drought is not inescapable. Based on these data, we calculated the relative leaf area (rLA) of each accession under both well-watered and drought conditions (see Materials & Methods; Fig. 2C; Table P1). We used rLA rather than the actual leaf area to allow comparison of the leaf area of each accession relative to the study population for each watering regime. We also show that the plasticity index derived from the difference in rLA between conditions (i.e. reflecting the slopes of the reaction norms in Fig. 2C; Table P1) is proportional to the fold change in leaf area under drought relative to well-watered conditions (Fig. S6).

**Fig. 2.**
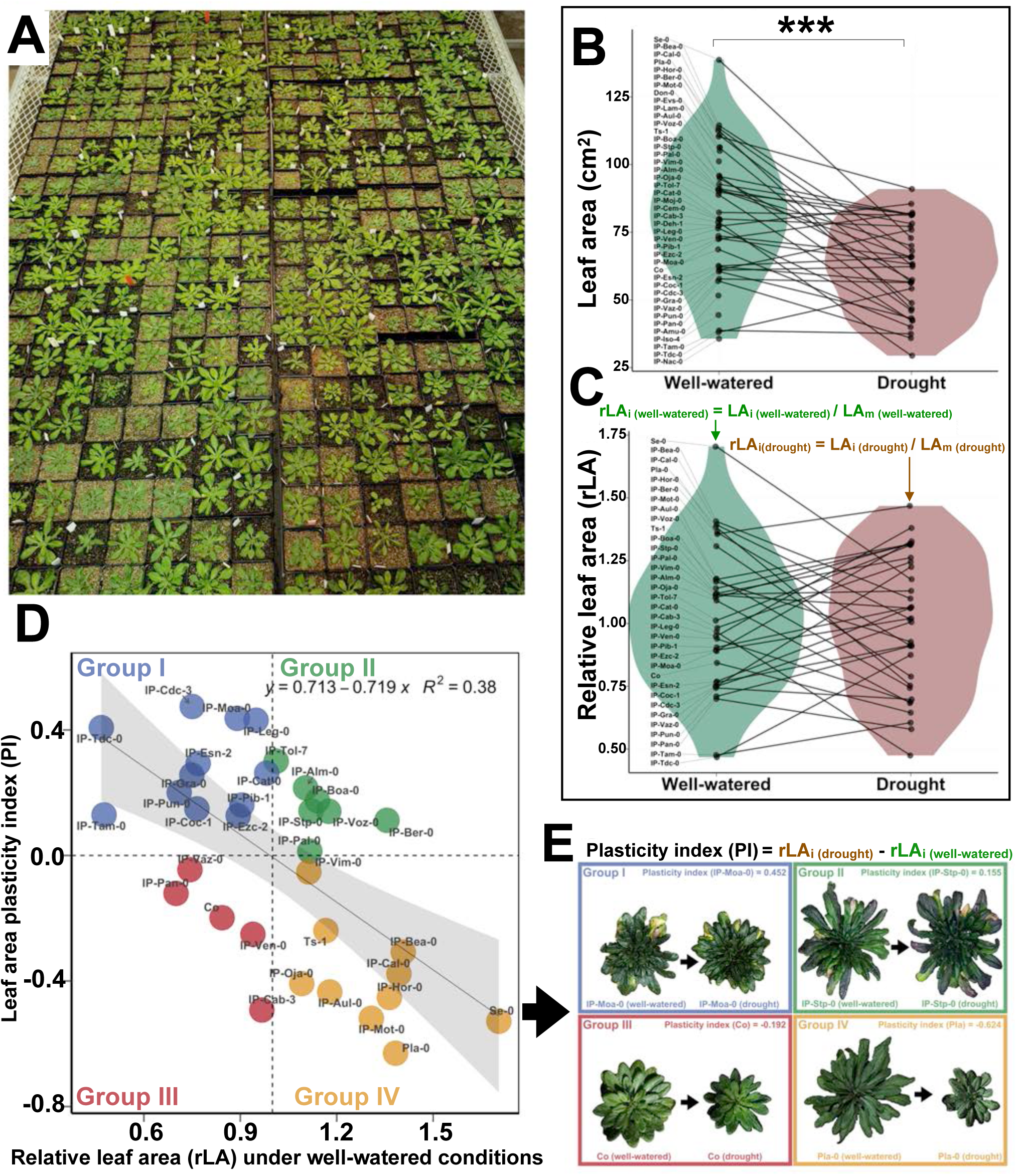
Accession-specific rosette leaf area responses in Arabidopsis. (A) Detail of the common garden experiment, illustrating the randomization of well-watered and drought treatments, as reflected in soil coloration. (B) Phenotypic reaction norms illustrating a reduction of leaf area in response to drought in most but not all accessions. Violin distributions represent probability densities of leaf area (cm^2^) among the accessions included in this study. Datapoints depict the mean phenotypic values of individual accessions, and the reaction norms represent the plastic response in the leaf area of each accession in response to drought (t-test; *P*-value < 0.001). (C) Phenotypic reaction norms illustrating plastic and homeostatic responses in relative leaf area (rLA). Violin distributions represent probability densities of rLA among the accessions included in this study. Datapoints depict the rLA of individual accessions, and the reaction norms represent the plastic response in rLA of each accession in response to drought (t-test; *P*-value > 0.05). (D) Negative relationship (*P*-value < 0.05) between leaf area plasticity index (slope of the reaction norms in 2C) in response to drought (y-axis) and rLA under well-watered conditions (x-axis). Data were fitted using a linear regression model. Grey shadow represents the 95% CI of the fit regression line. Four different quadrants are defined by two dashed lines: the average rLA (x-axis), and y = 0, which is the homeostatic point at which a change in water availability does not affect rLA (i.e. a plasticity index of 0). Data points depict individual accessions with their respective label. Data points are colored based on in which quadrant they fall (see text). (E) Images illustrating the plastic and homeostatic responses to drought found among the accessions included in this study.

We explored the relationship between rLA in the absence of drought and its plastic response (plasticity index; PI) to drought. Fig. 2D shows that in general accessions with higher rLA under well-watered conditions displayed greater plastic reductions in rLA when subjected to drought (negative leaf area plasticity; yellow datapoints in Figs 2D, E). However, this relationship was not invariant; we also observed individual accessions with high rLA under well-watered conditions and positive plasticity indexes under drought (green datapoints in Fig. 2D and Fig. 2E). Moreover a few accessions displayed low rLA and negative leaf area plasticity indexes (red datapoints in Fig. 2D and Fig. 2E), Most commonly, accessions with lower rLA under well-watered conditions displayed homeostatic phenotypic responses to drought (Fig. 2D, E blue datapoints).

While the severity of drought depends on multiple environmental parameters, its physiological response is very characteristic. It results in stomatal closure and a reduction in CO_2_ assimilation, as reflected in decreases in Net Primary Productivity (NPP), an index that, therefore, can be used as a reliable drought indicator (Huang *et al*., 2016). We determined that the plastic leaf area responses (negative values of leaf area plasticity index) we observed in our study were typical of accessions from more productive habitats as reflected in Net Primary Productivity in spring (NPP spring; Fig. 3A). In contrast, homeostatic leaf area responses (zero or positive values of leaf area plasticity index) were typical of accessions from less productive habitats (Fig. 3A).

**Fig. 3.**
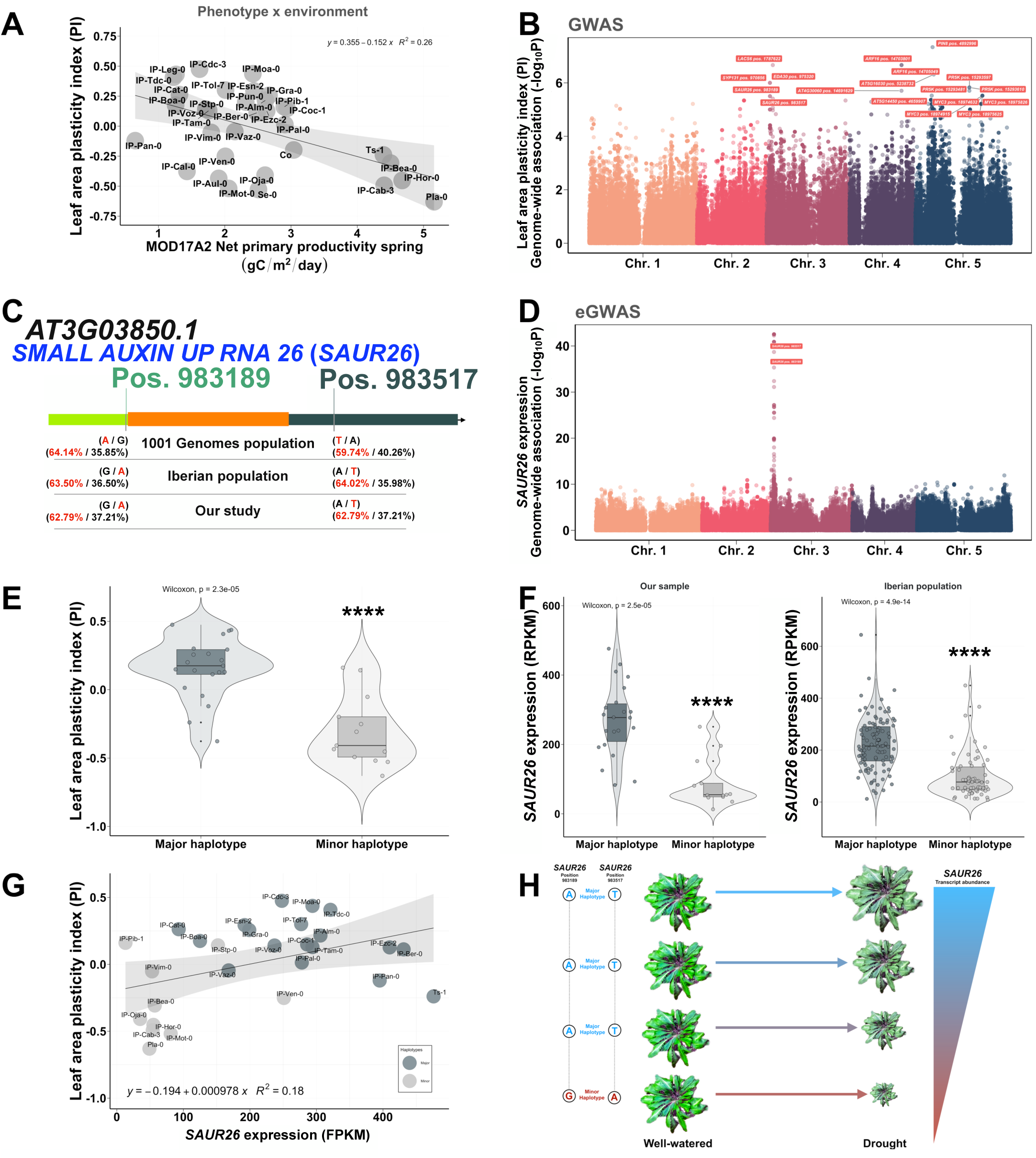
*SMALL AUXIN UP RNA 26* (*SAUR26,* AT3G03850) is a candidate regulator of Arabidopsis leaf area plasticity in response to drought. (A) Accessions from more productive areas display stronger negative reductions in leaf area plasticity relative to accessions from less productive environments. Scatter plot depicting the negative relationship (*P*-value < 0.05) between leaf area plasticity index (y-axis) and Net Primary Productivity during the spring season (x-axis). (B) Manhattan plot representing the strength of association between genetic variants and leaf area plasticity in response to drought. (C) *SAUR26* gene model, including the two co-varying SNPs highlighted in 2B. Orange block depicts the sole exon, green block the 5’ UTR, dark green block the 3’ UTR. Labels above indicate the position of the two SNPs that co-vary, forming a haplotype in our sample. Major/minor allele frequency in our sample, in the 189 accessions from the Iberian Peninsula included in the 1001 Genome Project, and the whole 1001 Genomes Project sample dataset is provided. (D) Manhattan plot representing the genome-wide strength of association between the genetic variation present within the 665 accessions from the transcript abundance reference panel (Kawakatsu *et al*., 2016) that overlap with the collection of accessions included in the 1001 Genomes Project, and the variation in the transcript abundance of *SAUR26* (Kawakatsu *et al*., 2016). The peak in chromosome 3 represents the cis-genetic variants in *SAUR26* affecting its transcript abundance. For the Manhattan plots in 3B and D, the score (y-axis) consists of the negative logarithm of the *P*-value and provides association strength. Each data point represents a SNP with a color corresponding to the chromosome where the genetic variants are located. The x-axis and the different colors shown in the SNPs depicted in the Manhattan plots represent the different chromosomes. The higher the score, the lower the *P*-value, and the stronger the association between genetic variation and leaf area plasticity or *SAUR26* transcript abundance. (E) In the Iberian population, the major haplotype of *SAUR26* is associated with homeostatic or positive responses of relative leaf area (rLA) in response to drought. Conversely, the minor haplotype in the Iberian population is associated with plastic responses in rLA in response to drought. Violin plots show significantly different probability densities of leaf area plasticity in response to drought for the major and minor allele haplotypes in *SAUR26.* (F) The haplotypes in *SAUR26* associated with leaf area plasticity/homeostasis in response to drought are associated with differences in *SAUR26* gene expression. Violin plots show significantly different probability densities of *SAUR26* transcript abundance for the major and minor allele haplotypes in *SAUR26* for both our sample and the entire Iberian population. (G) Scatter plot illustrating the positive relationship between leaf area plasticity index (y-axis) and *SAUR26* transcript abundance (x-axis). For 2A and 2G, the grey shadow represents the 95% CI of the fit regression line (dark grey). (H) Cartoon illustrating the regulatory role of genetic and transcript variation in *SAUR26* and the response of leaf area to drought.

Genetic variants significantly associated with leaf area plasticity were identified by GWAS (Fig. 3B; Table G3). Gene ontology (GO) analysis of functions associated with the genes harboring the variants identified by GWAS revealed an enrichment in auxin-related processes (Table GO1). Among the variants identified from GWAS (Fig. 3B; Table G3) as associated with leaf area plasticity, we highlight two co-varying SNPs in the 5’ and 3’ UTR that comprise a haplotype of *SMALL AUXIN UP RNA 26* (*SAUR26,* AT3G03850) (Fig. 3C). These variants were identified by GWA analysis using a linear model (Fig. 3B; Table G3). To confirm that the identification of these candidates was not an artifact due to population structure, we verified that these variants were also identified using two different analyses that correct for population structure in the sample: an Accelerated Mixed Model using GWAPP and an alternative Mixed Model approach, GWAS-Flow (these two SNPs are among the green datapoints in Fig. S7A and C; see Table G3).

We additionally found from TWA analysis that the transcript levels of *SAUR26* are also correlated with natural variation in leaf area plasticity in response to drought (Tables T1, T2). eGWAS confirmed the cis-regulatory effect of the same two genetic variants as depicted in Fig. 3C (cis-regulation; Fig. 3D; Table G6), consistent with the correlation between the genetic variation in *SAUR26*, its transcript abundance, and leaf area plasticity.

Fig. 3E illustrates the haplotypic differences in leaf area plasticity for *SAUR26* in our study. Accessions harboring the minor *SAUR26* haplotype display negative plasticity in rLA, while the major *SAUR26* haplotype is associated with homeostatic and positive plasticity responses in rLA (Fig. 3E; Wilcoxon *P*-value < 0.001). These haplotypic differences were also associated with the transcript levels of *SAUR26* (Fig. 3F), which we confirmed not only in the group of accessions included in our study (Wilcoxon *P*-value < 0.001) but also in the much larger group of 189 Iberian accessions included within the 1001 Genomes Project (Fig. 3F; Wilcoxon *P*-value < 0.001). Both the genetic variation defined by this haplotype in *SAUR26* and the resulting *SAUR26* transcript abundance are associated with the interplay between plasticity and homeostasis that we determine in our study (Figs 3E, F). These results highlight *SAUR26* as a candidate regulator of leaf area plasticity in response to drought (Figs 3G, H).

In our GWA analysis on rLA (Fig. S8A, Table G1, G2), no SNPs were associated with rLA under both well-watered and drought conditions, i.e., no SNPs were associated with homeostasis and thus horizontal lines are absent from Fig. S8A. The results from TWA analysis were strikingly different, identifying not only transcripts associated with leaf area plasticity but also transcripts associated with leaf area homeostasis (Fig. 4; Fig. S8B). Regarding homeostasis, four transcripts are positively correlated with rLA under both well-watered and drought conditions: *SOFL2 (AT1G68870), CPN60B (AT1G55490), AT5G39471, and CKL8 (AT5G43320)* (Fig. 4A). Accessions with higher transcript abundance of these genes display a higher rLA under both well-watered and drought conditions (Fig. 4B-E; note the similarity of the regression lines in each panel). Conversely, we identified four genes negatively correlated with rLA: *RHF1A* (AT4G14220), *AT5G60160*, *AT4G15260*, and *AT5G16800* (Fig. 4F). Accessions with a higher transcript abundance of these genes display lower rLA under both well-watered and drought conditions (Fig. 4G-J; note the similarity of the regression lines in each panel).

**Fig. 4.**
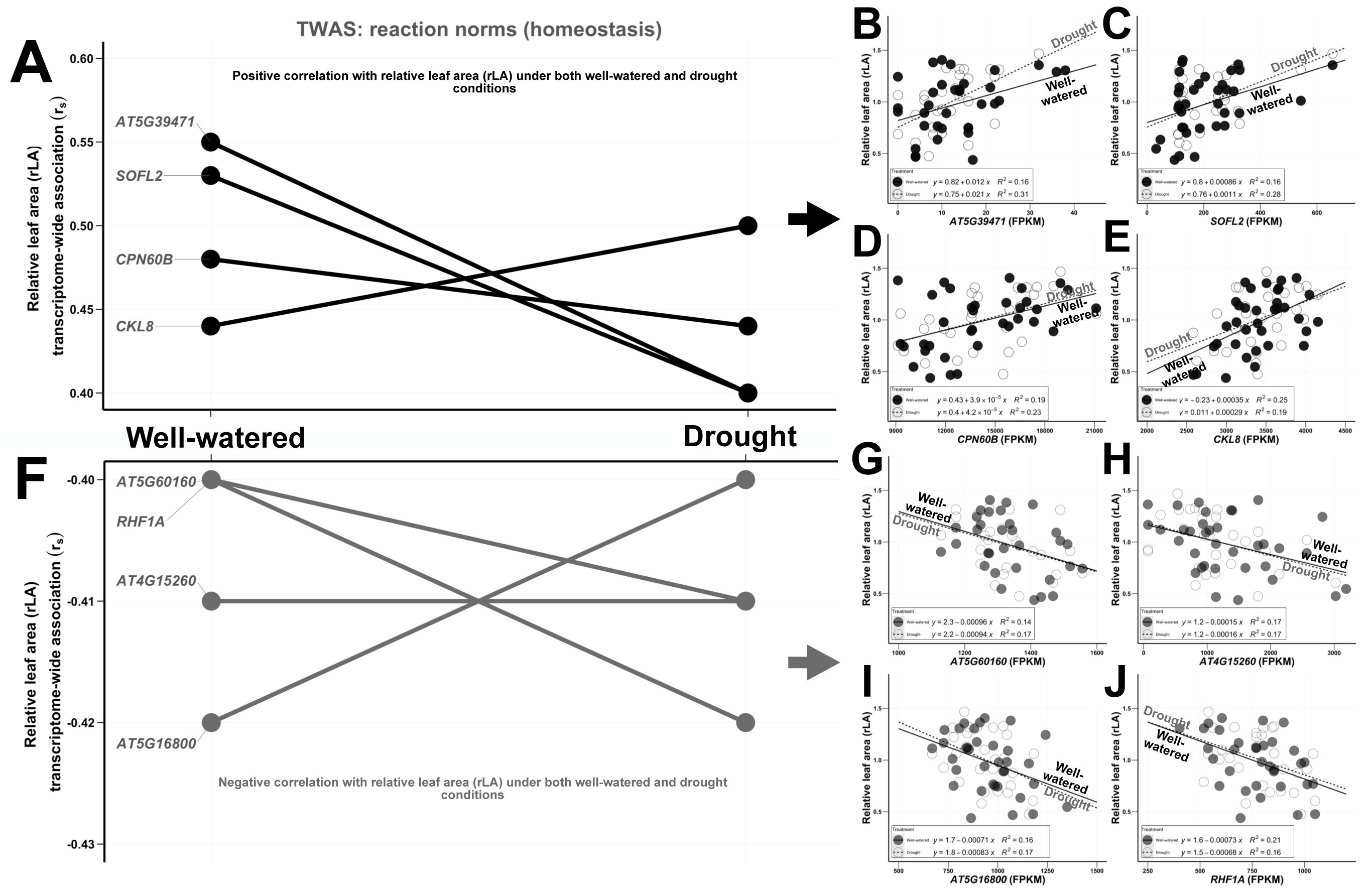
Transcriptomic reaction norms uncover genetic correlates of Arabidopsis leaf area homeostasis in response to drought. (A) homeostatic reaction norms for transcripts significantly and positively correlated with relative leaf area (rLA) under both watering regimes. (B) to (E). Scatter plots illustrating positive and homeostatic relationships (*P*-value < 0.05) between rLA under both well-watered (closed circles) and drought conditions (open circles) conditions (y-axis), and transcript abundance (FPKM; x-axis) of (B) AT5G39471, (C) *SOB FIVE-LIKE 2* (*SOFL2; AT1G68870*), (D) *CHAPERONIN 60 BETA* (*CPN60B*; *AT1G55490*), and (E) *CASEIN KINASE I-LIKE 8* (*CKL8*; *AT5G43320*). (F) homeostatic reaction norms for transcripts significantly and negatively correlated with rLA under both watering regimes. (G) to (J) Scatter plots illustrating negative and homeostatic relationships (*P*-value < 0.05) between rLA under both well-watered (closed circles) and drought conditions (open circles) (y-axis), and transcript abundance (FPKM; x-axis) of (G) AT5G60160, (H) *AT4G15260*, (I) *AT5G16800*, and (J) *RING-H2 GROUP F1A* (*RHF1A*; *AT4G14220*). For 3a and 3f, Spearman’s rank correlation coefficient (r_s;_ y-axis) indicates the correlation between the rLA and the natural variation in transcript abundance of each gene. In Figs B to E and G to J, data were fitted using a linear regression model. Also, in panels b-e and g-j, filled circles depict the rLA of accessions grown under well-watered conditions, while empty circles squares depict the rLA of accessions grown under drought.

### Trade-off between plasticity and intrinsic water use efficiency

It was not surprising to find a trade-off between growth and drought resistance among the accessions in our study (Fig. 2D), as this is a well-established phenomenon (Zhang *et al*., 2020). Accordingly, we hypothesized that those accessions that displayed a homeostatic response to drought minimized water loss, and thus would exhibit an increased intrinsic water-use efficiency (WUE_i_). Given that δ^13^C reflects intrinsic water use efficiency (WUE_i_) (Juenger *et al*., 2005; Easlon *et al*., 2014), we analyzed the δ^13^C values recorded by Dittberner *et al*. (2018), available for 21 of the accessions used in our study. The results confirm a trade-off between plasticity and WUE_i_ (Fig. 5A). Accessions that exhibit strong negative plasticity following drought (negative leaf area plasticity indices) have a lower WUE_i_ in the absence of drought (more negative δ^13^C values). Those accessions that maintain or increase leaf area in response to drought (positive leaf area plasticity indices) display a higher WUE_i_ in the absence of drought (more positive δ^13^C values).

**Fig. 5.**
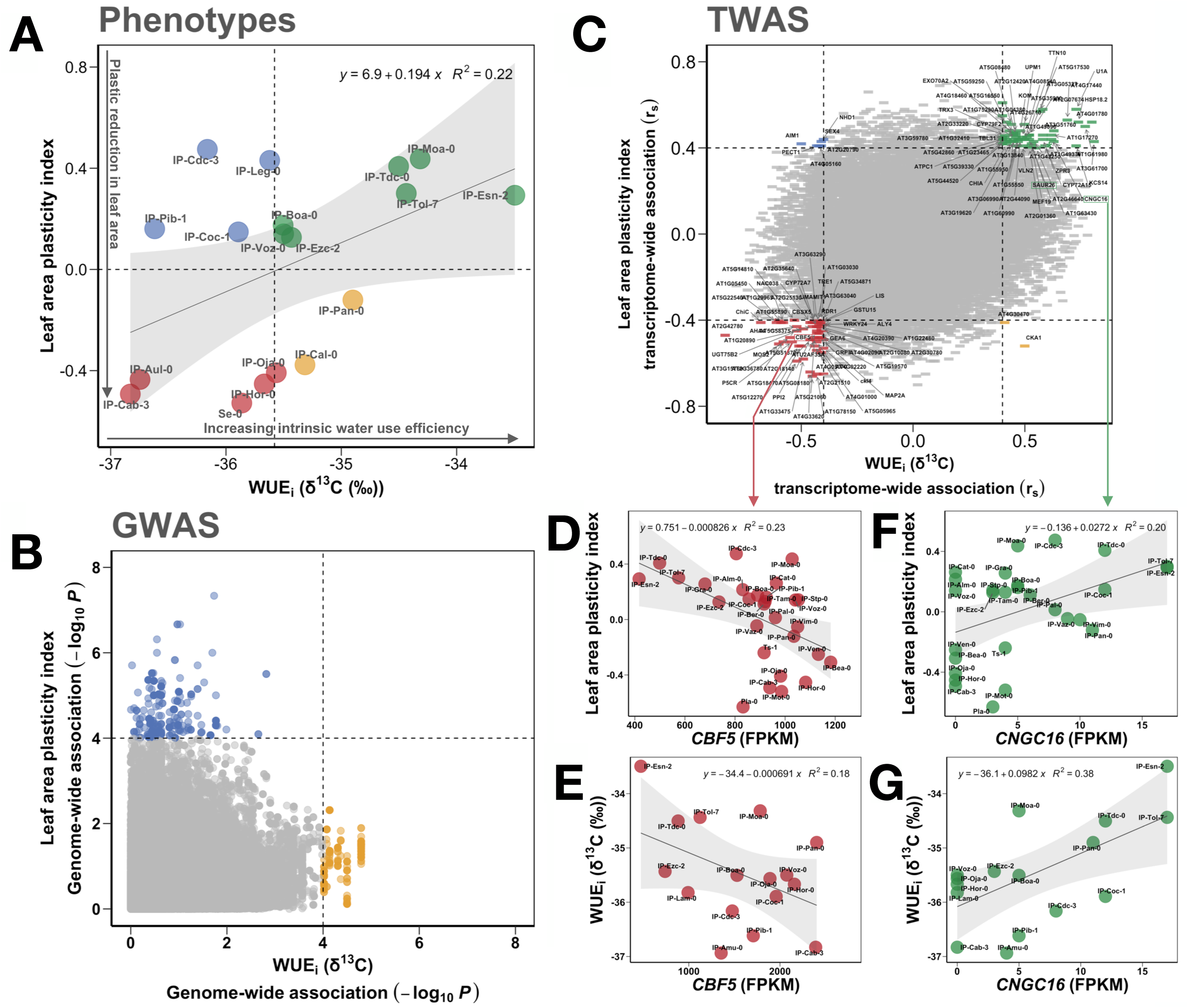
Arabidopsis accessions that display leaf area homeostasis in response to drought, display higher water use efficiency as defined by δ^13^C. (A) Scatter plot illustrating the positive relationship (*P*-value < 0.05) between leaf area homeostasis in response to drought (leaf area plasticity index, y-axis) and water use efficiency (x-axis) among Iberian Arabidopsis accessions used in this study. Data were fitted to a linear regression model. Grey shadow represents the 95% CI of the fit regression line (black). δ*^13^C* data for the accessions were retrieved from Dittberner *et al*. (2018). Four different quadrants are defined by two dashed lines: y = 0, which is the homeostatic point at which a change in water availability does not affect relative leaf area (i.e., a plasticity index of 0), and the average intrinsic water use efficiency (WUE_i_, δ^13^C; x-axis). Data points depict individual accessions with their respective labels. (B) The genetic basis of leaf area plasticity in response to drought and water use efficiency. This plot depicts the respective relationship between the GWAS associations found for the leaf area plasticity index in response to drought (y-axis) and those for WUE_i_ (Dittberner *et al*., 2018) (x-axis). Data points represent individual SNPs. The y-axis and x-axis depict the extent of the association with, respectively, leaf area plasticity, and WUE_i_, as the negative logarithm of the p-value. Dashed lines depict an arbitrary threshold for significance that is consistent across comparisons. (C) Transcriptomic association with plastic leaf area responses to drought and WUE_i_. Spearman’s rank correlation depicts the relationship between the association of the transcript abundance of each gene in the Arabidopsis genome with leaf area plasticity in response to drought (r_s_) on the y-axis and with WUE_i_ on the x-axis. Dashed lines define an arbitrary threshold to consider transcripts correlated positively (r_s_ ≥ 0.4) or negatively (r_s_ ≤ −0.4) associated with leaf area plasticity (y-axis) or WUE_i_ (x-axis). *CBF5* and *CNGC16* exemplify transcripts with opposing relationships to leaf area plasticity and water use efficiency as shown in 4C, D and E. Scatter plots depicting the relationship (P-value < 0.05) between leaf area plasticity index in response to drought (4D), or water use efficiency (4E) on the y-axis, and transcript abundance (FPKM) of *CBF5* on the x-axis. F and G. Scatter plots depicting the relationship (*P*-value < 0.05) between (4F) leaf area plasticity index in response to drought or (4G) water use efficiency (Dittberner *et al*., 2018) on the y-axis, and transcript abundance (FPKM) of *CYCLIC NUCLEOTIDE-GATED CHANNEL 16* (*CNGC16*) on the x-axis. In all panels, data were fitted using a linear regression model. Grey shadow represents the 95% CI of the fit regression line.

GWA analysis failed to identify any genetic variants associated with the observed phenotypic trade-off between intrinsic WUE_i_ and leaf area plasticity (Fig. 5B; Table G3, G4), i.e., no datapoints in the upper right quadrant of Fig. 5B. However, TWA analysis (Fig. 5C) provided insight into this trade-off. Red datapoints in Fig. 5C are transcripts with greater abundance in those negatively plastic and less water-use efficient accessions depicted in red in Fig. 5A; GO analysis revealed that these were enriched in processes involved in RNA modification (Table GO2); we chose the putative pseudouridine synthase *CBF5* (*AT3G57150*) as an example (Figs 5D, E). Conversely, green points in Fig. 5C represent transcripts with greater abundance in the homeostatic and water-use efficient accessions depicted in green in Fig. 2D. We chose *CYCLIC NUCLEOTIDE-GATED CHANNEL 16* (*AT3G48010*; *CNGC16*) as an example (Figs 5F, G).

We determined that in our study, accessions that in response to drought displayed either increased or homeostatic leaf area plasticity also had higher WUE_i_ in the absence of drought (Fig. 5A). We then conducted a meta-analysis on the 95 Iberian accessions with published δ^13^C information (Dittberner *et al*., 2018). GWA analysis of these accessions rendered a distinctive peak in a single gene, identifying a haplotype consisting of eight co-varying SNPs in *ClpX3* (*AT1G33360*; Fig. 6A; Table G5), an ATP-dependent Clp protease subunit that plays essential roles in chloroplast development (Kim *et al*., 2009). Consistent with this result but more informative was the TWA analysis of WUE_i_ in the 88 Iberian accessions with both published transcript abundance and δ^13^C information (Fig. 6B): we uncovered a distinctive enrichment in photosynthesis-related genes (Fig. 6C; Table GO3) among transcripts that are upregulated in accessions with high WUE_i_ (more positive δ^13^C values; Fig. 6B; Table T1, T2). We highlight the positive relationships with WUE_i_ uncovered for the transcript abundance of *RuBisCO small subunit 2B* (*AT5G38420*; *RBCS2B;* Fig. 6D) and the positive relationship between the transcript abundance of *SAUR26* and WUE_i_ (Fig. 6E), in concordance with other findings in our study.

**Fig. 6.**
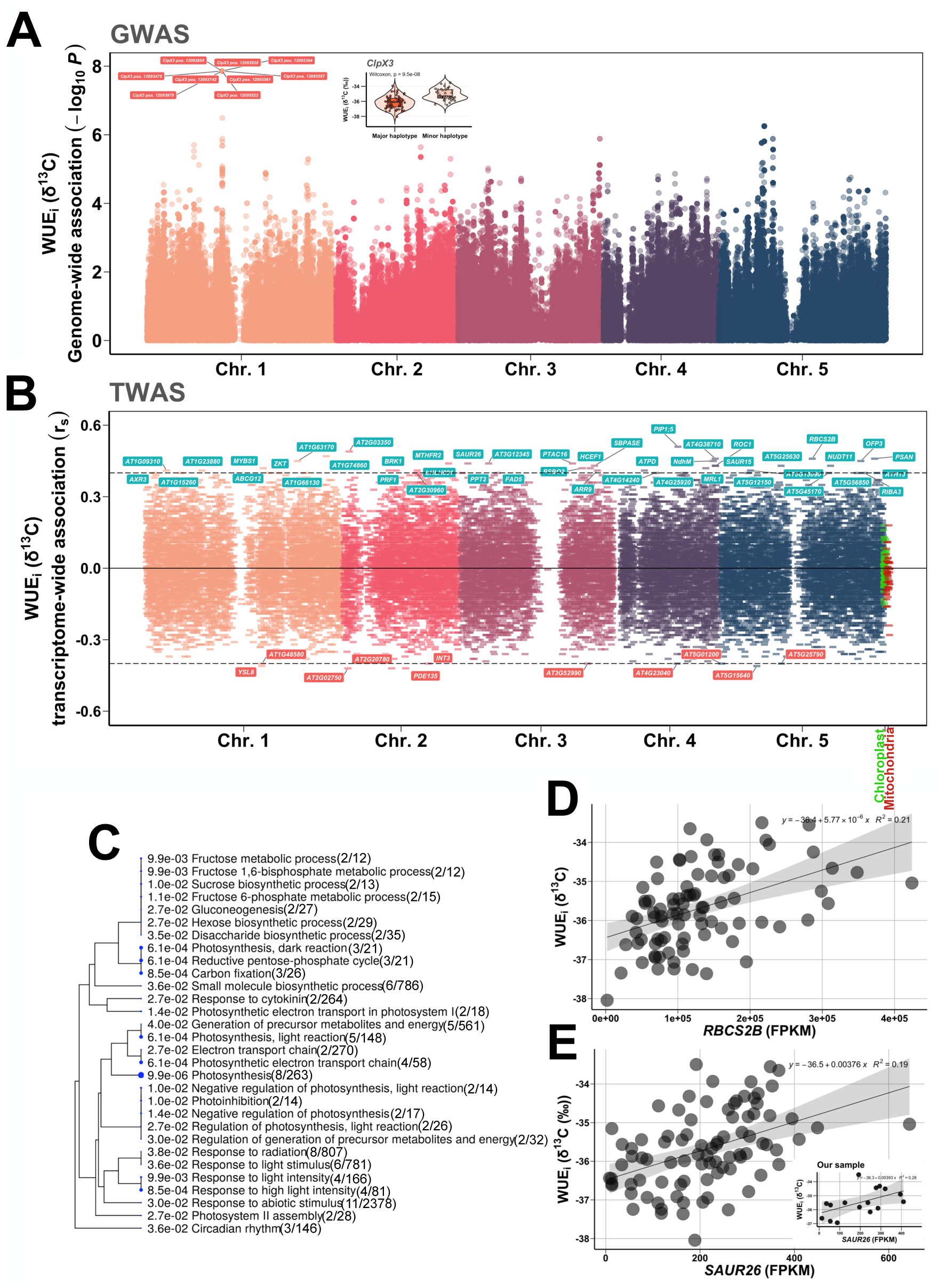
Photosynthesis-related transcript variants regulate water use efficiency in Iberian Arabidopsis accessions**. (**A) Manhattan plot representing the genome-wide strength of association between genetic variation and variation in δ^13^C present within 95 Iberian accessions with δ^13^C values available from Dittberner *et al*. (2018). The score (y-axis) consists of the negative logarithm of the *P*-value and provides association strength. Inset: Violin plots representing the probability densities of WUE_i_ for the major and minor allele haplotypes in *ClpX* for the accessions included in Fig. 6A. (B) T-Manhattan plot representing transcriptome-wide strength of association between variation in transcript abundance (Kawakatsu *et al*., 2016) and natural variation in δ^13^C. This analysis was performed on all Iberian accessions with both transcript abundance data (Kawakatsu *et al*., 2016) and δ^13^C information (Dittberner *et al*., 2018) (n = 88 accessions). The relationship is defined by the Spearman’s rank correlation coefficient (r_s_). Dashed lines define an arbitrary threshold for positive (r_s_ ≥ 0.4) or negative (r_s_ ≤ −0.4) correlation with water-use efficiency (y-axis). Blue labels highlight the transcripts positively associated with higher WUE_i_, while red labels highlight those negatively associated with higher WUE_i_. (C) Hierarchical clustering tree summarizing the correlation among significant GO Biological Process categories resulting from analysis of the positively correlated candidates resulting from TWA analysis of water use efficiency determined from δ^13^C values in 88 Iberian Arabidopsis accessions. Ratios provide the number of genes positively correlated with WUE_i_ within each GO category relative to the total number of genes within that category. The size of the blue dots increases proportionally with P-value significance. (D) Scatter plot depicting the positive relationship (*P*-value < 0.05) between WUE_i_ (y-axis) and *RuBisCO SMALL SUBUNIT 2B* (*AT5G38420*; *RBCS2B*) transcript abundance (x-axis) for the Iberian accessions included in Fig. 6B. (E) Scatter plot depicting the positive relationship (*P*-value < 0.05) between intrinsic water-use efficiency (WUE_i_; y-axis) and *SMALL AUXIN UP RNA 26* (*SAUR26,* AT3G03850) transcript abundance (x-axis) for the Iberian accessions included in 6b. Inset: Scatter plot depicting the positive relationship (*P*-value < 0.05) between intrinsic water-use efficiency (WUE_i_; y-axis) and *SMALL AUXIN UP RNA 26* (*SAUR26,* AT3G03850) transcript abundance (x-axis) for the 43 accessions included in our study. For 5D, and 5E, gray shadow represents the 95% CI of the fit regression line (black).

## DISCUSSION

### Identifying candidate regulators of plasticity and homeostasis

An adaptive trade-off between growth and drought resistance in plants is well-established (Zhang *et al*., 2020). In productive environments, accessions with higher rLA in the absence of drought may have an advantage in competing for aboveground resources relative to smaller individuals, as proposed by Tilman (1988). On the other hand, plants adapted to unproductive environments typically fix less carbon, have higher WUE_i_, and display reduced vegetative growth, even when grown in rich environments (Parsons, 1968; Chapin, 1991); however, these plants are typically stress-tolerant (Billings and Mooney, 1968; Grime, 1974, 1977). In our study, accessions typical of more productive areas display strong and negative plastic responses to drought (Fig. 3A). Conversely, accessions with homeostatic or even positive plasticity in rLA in response to drought typify less productive areas (Fig. 3A). Here, we sought to identify the genetic basis of these contrasting ecological strategies.

Our main interest was not to identify natural variation associated with well-watered or drought phenotypes, but rather natural variation associated with the degree of plasticity evoked in response to drought, i.e., natural variation associated with reaction norms, rather than with phenotypes in one environment or the other (well-watered or droughted). In exploring the genetic basis of plasticity, we identified 31 genes corresponding to the top 50 candidate SNPs with the strongest association with the leaf area plasticity index (Fig. 3B; Table 1; Table G3). To the best of our knowledge, 20 out of these 31 genes have been previously characterized, and many of these are related to auxin regulation and leaf expansion. This enrichment in auxin-related processes was confirmed by GO analysis (Table GO1). Notably, when we compare the candidates resulting from GWA analysis on rLA vs. leaf area plasticity, only eight candidate SNPs overlapped between both traits (green datapoints in Fig. S9A), and none of these eight SNPs are in genes related to auxin. In other words, the auxin-related candidate genes are related to *plasticity* in this trait and not to the trait itself.

**Table 1.**
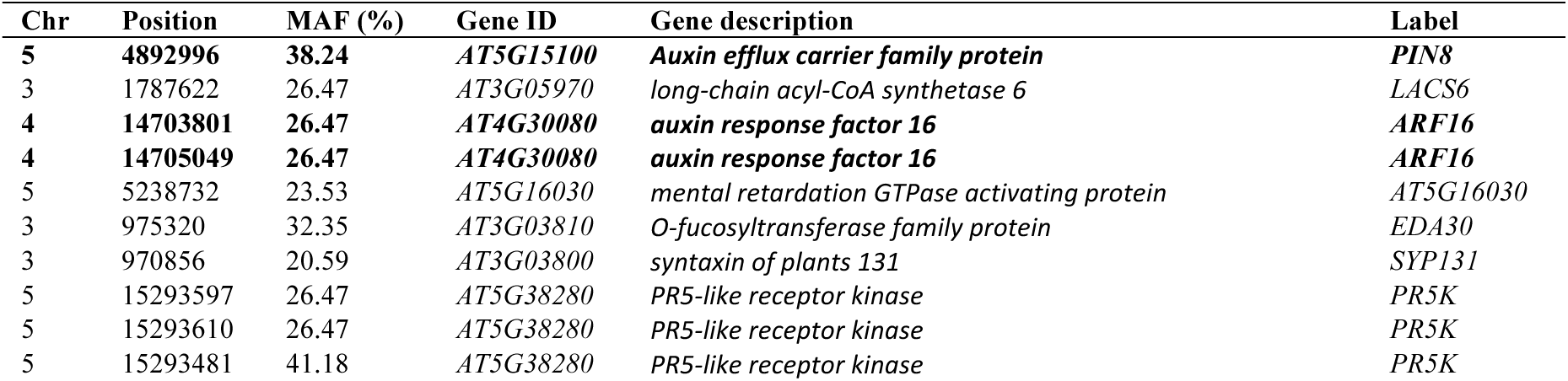
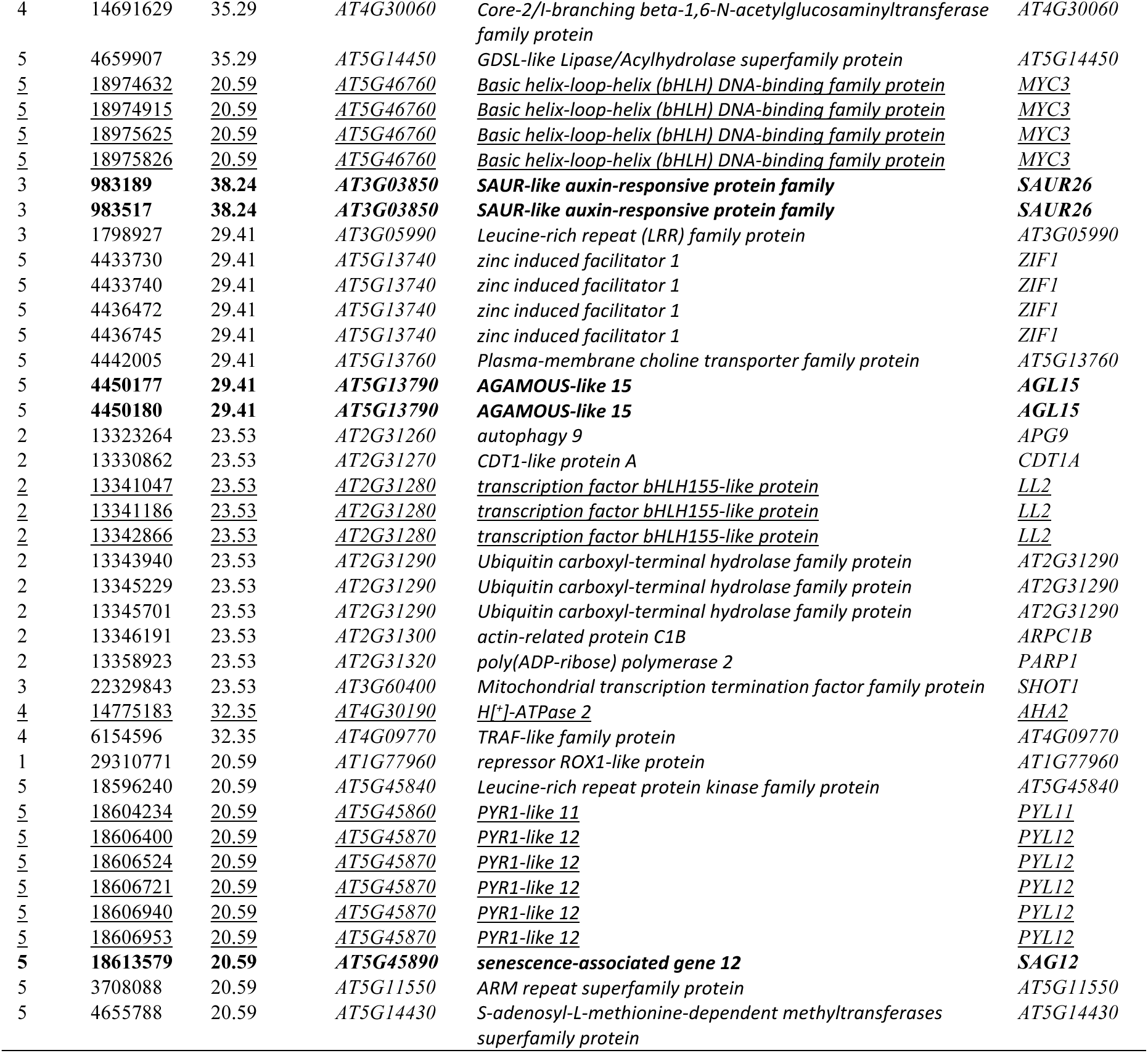
List of 31 genes corresponding to the top 50 candidate SNPs with the strongest association with the leaf area plasticity index. Genes encoding proteins with auxin-related functions from GO analysis are bolded; additional genes related to auxin based on literature analysis are underlined.

The variant with the strongest association with the plastic response to drought was the auxin efflux carrier family protein *PIN8* (*AT5G15100*). Among the associated SNPs, we also found two co-varying SNPs in *AUXIN RESPONSE FACTOR 16* (*ARF16*; *AT4G30080),* two co-varying SNPs in *SMALL AUXIN UP RNA 26* (*SAUR26*; *AT3G03850*); two variants in *AGAMOUS-like 15* (*AT5G13790*; *AGL15*), a gene that negatively regulates auxin signaling in Arabidopsis (Zheng *et al*., 2016); and three variants in *LONESOME HIGHWAY LIKE 1* (*LL2*; *AT2G31280*), an atypical basic helix-loop-helix (bHLH) transcription factor that regulates early xylem development downstream of auxin (Ohashi-Ito *et al*., 2013).

Among the candidates that are not formally included in the GO category “response to auxin,” we found six co-varying SNPs in the abscisic acid receptors *PYR1-like 11* (*PYL11*; *AT5G45860*) and *PYR1-like 12* (*PYL12*; *AT5G45870*). ABA and auxin pathways interact in the regulation of growth (Emenecker & Strader 2020), and also in the regulation of plasma membrane H^+^ ATPases (of which there is one member in the list, *H^+^ ATPase 2* (*AHA2; AT4G30190*)) during cell expansion (Spartz *et al.,* 2014) and stomatal control (Wong *et al*., 2021). We also found four co-varying SNPs in the *Basic helix-loop-helix (bHLH) DNA-binding family protein MYC3* (*AT5G46760*), a known regulator of glucosinolate biosynthesis (Schweizer *et al*., 2013). Auxin negatively regulates glucosinolate levels, which appears to integrate growth and stomatal regulation during drought (Salehin *et al*., 2019) and ABA response (Zhao *et al*., 2008). GO biological process analysis on positively correlated candidates resulting from TWAS analysis on leaf area plasticity index for drought also revealed an enrichment of glucosinolate biosynthesis genes (Table GO4).

We found that among the auxin-related variants associated with leaf area plasticity (Fig. 3B), SNPs in *SAUR26* were of particular interest. The majority of characterized *SAUR* genes are positive regulators of plant growth (Spartz *et al*., 2014; Li *et al*., 2015; Stortenbeker and Bemer, 2019). Previous studies identified a regulatory role of *SAUR26* in rosette growth plasticity in response to temperature changes (Wang *et al*., 2019; Wang *et al.,* 2021). High temperatures accelerate evaporative water loss from the plant while drought limits water availability, and these stresses commonly co-occur. A reduction in leaf area is a common adaptive mechanism to maintain the relative water content of plants under water deficit (Anyia and Herzog 2004). These previous studies serve as validation of the role of *SAUR26* in regulating rosette plasticity to drought stress as implicated by our study (Figure 2).

The two variants in *SAUR26* associated with rosette size were found in the 5’ and 3’ UTR, respectively (Fig. 3C), regions of the transcript that typically regulate gene expression. Indeed, we found the transcript levels of *SAUR26* to be associated with this genetic variation in *SAUR26* itself (cis-regulation; Fig. 3D), which we confirmed both in the group of accessions included in this study and, importantly, in the larger group of 189 Iberian accessions in the 1001 Genomes Project (Fig. 3F). The minor haplotype is associated with negative plasticity in leaf area in response to drought. Conversely, the major haplotype, associated with higher transcript abundance of *SAUR26*, is associated with leaf area homeostasis and positive plastic responses (Fig. 3E).

The location of these *SAUR26* variants in non-coding regions of the mRNA implicated alteration in transcript abundance as a molecular phenomenon underlying the phenotypic correlations. Indeed, natural variation in transcript abundance of *SAUR26* explained 18% of the observed plastic response in leaf area to drought in our study (Fig. 3G, H). The importance of *SAUR26* transcript abundance is supported by previous work that identified cis-elements in the 5’ upstream region or the 3’UTR region of *SAUR26* that were associated with growth thermo-responsiveness of the rosette compactness index (Wang *et al*., 2019), (Wang *et al*. 2021), again validating our work. This conclusion is also supported by our meta-analysis of the larger Iberian population with transcriptome data, which also identified *SAUR26* transcript abundance as positively correlated with higher WUE_i_ (Fig. 6B, E), along with other related auxin-related transcripts such as *AUXIN RESISTANT 3* (*AXR3*, also known as *IAA17*, *AT1G04250;* Fig. S10A) (Leyser *et al*., 1996) and *SMALL AUXIN UPREGULATED 15* (*SAUR15*, *AT4G38850*; Fig. S10B) (Oh *et al*., 2014). As we uncover in this study, both transcript abundance and natural genetic variation in *SAUR26* cis-regulatory elements are associated with the interplay between plasticity and homeostasis in rLA following drought stress (Fig. 3, Tables G3, T1, T2).

Interestingly, the minor haplotype frequency for this *SAUR26* haplotypic variant in our study sample (minor haplotypic frequency, MHF = 38.09%) is similar to the frequencies we find in the 189 accessions within the Iberian population (MHF = 35.98%). However, the minor haplotype in the Iberian population becomes the haplotype in higher frequency (64.14%) in the global 1001 Genomes collection (The 1001 Genomes Consortium, 2016). It has been shown that Iberian Arabidopsis populations locally adapted to drought conditions in this region out-perform accessions from wetter areas of Northern Europe when both are exposed to drought (Exposito-Alonso *et al*., 2019). Following this, given that the major haplotype in our study is associated with homeostatic responses to drought (Fig. 3E) and that its frequency becomes reduced outside the drought-prone Iberian region, we can speculate that this *SAUR26* haplotypic variant is under selection in the Iberian population, conferring an adaptive advantage to the water-limited conditions of the region.

We did not identify any SNPs associated with rLA under both well-watered and drought conditions. Therefore, the genetic basis of growth homeostasis under drought could not be defined in our study using GWAS (Fig. S8A). On the other hand, the reaction norms based on the strength of association of transcript abundance with rLA, i.e., TWAS, identify four transcripts that are positively correlated with rLA under both well-watered and drought conditions (Figs 4A-E, S8B). This means that accessions with greater transcript abundance for those genes display higher rLA under both well-watered and drought conditions. Conversely, we identified four transcripts negatively correlated with rLA such that accessions with a higher transcript abundance of those genes display lower rLA under both well-watered and drought conditions (Fig. 4F-J). Therefore, these eight genes are strong functional candidates for the homeostatic regulation of rLA.

Of the eight resultant candidate genes, *CPN60B* (homeostatic, higher rLA) encodes a rubisco large subunit-binding protein (Cheng *et al*., 2020). *SOFL2* (homeostatic, higher rLA) is implicated in the control of cytokinin content (Zhang *et al*., 2009), which can contribute to drought tolerance (Rivero *et al*., 2007), while *AT4G15260* (homeostatic, lower rLA) is a putative UDP-glycosyltransferase, with another member of this superfamily identified as a cytokinin glycosyltransferase (Hou *et al*., 2004) that improves vegetative-stage drought survival when overexpressed. *AT5G16800* encodes a N-terminal acetyl transferase for which null mutants exhibit sensitivity to salinity stress. We can surmise functions in homeostatic control of leaf area for these four genes based on their described functions and phenotypes. The other four homeostatic genes are *AT5G39471* (annotated as a “Cysteine/Histidine-rich C1 domain family protein”) and *CKL8* (annotated as a casein kinase), both correlated with homeostatic and high rLAs, and *RHF1A* (RING-finger E3 ubiquitin ligase (Liu *et al*., 2008) and *AT5G60180* (vacuolar aminopeptidase (Park *et al*., 2017), both correlated with homeostatic and low rLAs. For these genes further research is needed regarding how they might contribute to homeostasis.

### Trade-off between relative leaf area (rLA), plasticity and WUEi

Plant species with low relative growth rates exhibit high WUE_i_ (Angert *et al*., 2009; Kimball *et al*., 2012). We can hypothesize that those plants with lower growth rates will have better drought resistance, consequently displaying growth homeostasis. These observations are both consistent with results from ecological studies and confirmed in crop species (Kasuga *et al*., 1999; Ferrero-Serrano and Assmann, 2016; Ferrero-Serrano *et al*., 2018; Ferrero-Serrano et al., 2024). In our study, accessions with lower rLA and homeostatic responses to drought display a greater abundance of transcripts encoding functions related to ion transport (yellow datapoints in Fig. S9B; Tables T1, T2, GO5), which could be related to osmoregulation and stomatal control of transpiration. Consistent with this argument, the homeostatic response in rLA in response to prolonged drought that we observe in our study correlates positively with a high predicted WUE_i_ recorded by Dittberner *et al*. (2018) for the same accessions (Fig. 5A).

We then studied the genetic basis of this trade-off between rLA, plasticity, and WUE_i_. There were no SNPs associated with both the leaf area plasticity index and WUE_i_ (Fig. 5B). However, when we compare the candidates obtained from TWA analysis, we identify transcripts associated with both low WUE_i_ and a negative plastic response in leaf area following sustained drought (Fig. 5C). Conversely, we also identify a set of transcripts that is significantly associated with both leaf area homeostasis (more positive leaf area plasticity index) and increased WUE_i_ (Fig. 5C). To exemplify these opposed responses to drought, given the enrichment in RNA modification and ion transport genes that we found in the transcripts regulating the trade-off between relative leaf area and plasticity (Fig S12B; Tables T1, T2), we highlight two of the genes identified by TWAS, with opposite correlations between transcript abundance and relative leaf area plasticity (Figs 5D, E vs. Figs 5F, G): (*CBF5*; *AT3G57150*) and *CYCLIC NUCLEOTIDE GATED CHANNEL 16* (*CNGC16*; *AT3G48010*).

*CBF5* encodes a protein with homology to a category of pseudouridine synthases, and its knockout in Arabidopsis is lethal (Lemontova *et al.,* 2007). Originally only non-coding RNAs, particularly rRNAs, were thought to be pseudouridylated, but it is now recognized that mRNAs are also a target for pseudouridylation. In both yeast and human cells, this post-transcriptional modification is induced by heat shock, with consequent impacts on splicing, mRNA stability, and translation (Karijolich *et al*., 2015), and some candidate pseudouridine synthases in rice have been shown to be transcriptionally regulated by drought (Dingra et al., 2023). We observe that accessions with higher transcript abundance of *CBF5* display both strong negative plastic responses in rLA in response to drought (Fig. 5D) and reduced WUE_i_ (Fig. 5E). Our observations suggest that stress-induced pseudouridylation may also occur in plants, with functional consequences for drought tolerance; other types of mRNA modifications in plants have been correlated with increased mRNA stability and translation under salinity stress (Kramer et al., 2020) and improved heat tolerance (Tang *et al.,* 2020).

Conversely, transcript abundance of *CNGC16* is associated with homeostatic responses in rLA and increased WUE_i_. CNGCs are a superfamily of ion channels permeable to divalent and monovalent cations (Köhler *et al*., 1999; DeFalco *et al*., 2016). *CNGC*s like *CNGC5* and *CNGC6* have been reported to be the basis of a cyclic GMP (cGMP)-activated nonselective Ca^2+^-permeable cation channel activity in the plasma membrane of Arabidopsis guard cells (Wang *et al*., 2013) associated with stomatal closure. *CNGC16* is also critical for reproductive success following drought stress (Tunc-Ozdemir *et al*., 2013). Consistent with these functional roles, in our study, higher transcript abundance of *CNGC16* is found in accessions with positive leaf area plasticity indexes under drought (leaf area homeostasis; Fig. 5F); and with high WUE_i_ (Fig. 5G).

Given the increased plastic responses in leaf area found in accessions with lower WUE_i_, we next conducted a meta-analysis on the larger set of 95 Iberian accessions with δ^13^C information from a previous study focused on the global Arabidopsis population (Dittberner *et al*., 2018). We identified a distinctive peak consisting of eight co-varying SNPs in a single gene, *ClpX3* (*AT1G33360*; Fig. 6A), an ATP-dependent Clp protease subunit (Nishimura and van Wijk, 2015). Null mutants in a different chloroplast protease subunit, ClpR, display a remarkable loss of chlorophyll (95 to 98% reduction) and photosynthetic capacity, turning entirely white when grown under high light (Kim *et al*., 2009).

TWA analysis of WUE_i_ determined from δ^13^C values in the 88 Iberian accessions with both δ^13^C information (Dittberner *et al*., 2018) and published transcript abundance (Kawakatsu *et al*., 2016) further supported the relationship with photosynthesis. Our TWAS on WUE_i_ in the larger Iberian population identified 31 out of the 45 candidate genes for WUE_i_ that we also identified in the subset of 20 Iberian accessions in our experimental study for which published δ^13^C data were available (Table T1 and T2). Gene Ontology (GO) biological process analysis of the resultant 45 candidates positively correlated with δ^13^C uncovered an unequivocal enrichment in photosynthesis-related genes for transcripts that are upregulated in accessions with high WUE_i_ (less negative δ^13^C values; Figs 6B, 6C; Table GO3). Among the candidates from our TWA analysis that are positively related to WUE_i_ was *RuBisCO small subunit 2B* (*AT5G38420*; *RBCS2B;* Figs 6B, D). An upregulation in *RuBisCO* may compensate for suboptimal CO_2_ concentrations at the sites of carboxylation arising from reduced stomatal apertures in plants from drought-prone environments. We illustrate eight additional genes with transcript abundances that were positively associated with WUE_i_ in Fig. S10. Among these are *NADH dehydrogenase-like complex M* (*NdhM*, *AT4G37925;* Fig. S10C), required for *NdH* complex assembly (Ifuku *et al*., 2011), and *High cyclic electron flow 1* (*HCEF1*; *AT3G54050*; Fig. S10D; Livingston *et al*., 2010), both regulators of cyclic electron flow around PSI (Yamori *et al*., 2011), *ATP-Synthase δ-Subunit* (*ATPD*, *AT4G09650*; Fig. S10E) (Maiwald *et al*., 2003), *Photosystem I reaction center subunit PSI-N* (*PSAN*, *AT5G64040;* Fig. S10F) (Haldrup *et al*., 1999), and *Photosystem II subunit O-2* (*PSBO2*; *AT3G50820;* Fig. S10G) (Murakami *et al*., 2005). Also, the candidate with the highest association score is a member of the plasma membrane intrinsic protein (PIP) subfamily of aquaporins (*PIP1;5*; *AT4G23400*; Fig. S10H). PIPs regulate water flux in guard cells as well as membrane permeability to CO_2_ in the mesophyll, with an important effect on mesophyll conductance and CO_2_ concentration at the sites of carboxylation (C_c_) (Groszmann *et al*., 2017; Zait, Ferrero-Serrano & Assmann 2021). Thus, *PIP1;5* is also a candidate regulator of photosynthetic rates. As discussed before, *SAUR26* is also among these candidates (Fig. 6B, E), increasing our confidence in the role of this gene as a regulator of plastic responses to drought (Fig. 3).

Among the candidates that are negatively related to high WUE_i_ there is an enrichment in carbohydrate transport-related processes (Table GO6). We can speculate that this enrichment reflects the transport of carbohydrates produced during photosynthesis to promote plant growth. This would be concordant with the trade-off between growth and WUE_i_ that we observe in our study.

## Conclusions

The adaptive role of plasticity has been at the center of a long-standing debate. Much work and discussion have focused on the principle that if plasticity is indeed adaptive, it may be under genetic control (Bradshaw, 1965; Via *et al*., 1995); however, few relevant genes have been identified. Here, we propose that not only plasticity but also homeostasis is under genetic control and subject to natural selection. This concept is supported by our dissection of the genetic basis of both plasticity and homeostasis in leaf area following drought, which identifies specific genes implicated in each process.

Knowledge from natural populations of how plants adapt to their environment can be incorporated in breeding and biotechnological approaches to crop improvement. In this study, we observe a trade-off between growth and drought response. A similar trade-off between stress tolerance and productivity threatens food security. It is noteworthy that we also found accessions that did not display this trade-off, e.g., those accessions marked in green in Fig. 2D, E. These accessions may be of particular interest from an agronomic perspective.

While candidate genes other than *SAUR26* identified in our study await experimental validation, their identification represents a significant step forward in understanding the genetic basis and mechanisms of plasticity and homeostasis. In particular, the concerted use of TWAS and GWAS increases the power to link genes to phenotypes (Kremling et al., 2019). A similarly integrative approach has identified water use efficiency traits in sorghum (Ferguson *et al*., 2020). Here, this approach has yielded insights into the genetic basis of response to resource limitation, identifying both candidate “plasticity genes” and candidate “homeostasis genes” for further study. In particular, the identification of potential homeostatic mechanisms has important implications for maintaining crop yield in the face of climate change. Results arising from diverse genome-wide analyses in both model and crop species (Schindele *et al*., 2020) can now be used to inform powerful new CRISPR-based approaches for gene editing and modulation of transcript abundance toward crop improvement (Abudayyeh *et al*., 2017).

## Supporting information

The following supplemental materials are available:

**Fig. S1.** Rosette leaf area observations at the time of flowering (Table P1).

**Fig. S2.** Quantile-quantile (QQ) plots for GWAS analysis using the Linear Model.

**Fig. S3.** Quantile-quantile (QQ) plots for GWAS analysis using the Accelerated Mixed Model.

**Fig. S4.** Quantile-quantile (QQ) plots for GWAS analysis using GWAS-Flow.

**Fig. S5.** Histograms of the distribution of r_s_ and scores obtained from GWA and TWA analyses of the traits discussed in this study illustrate the stringent significance thresholds imposed in this study (vertical dashed lines).

**Fig. S6.** The plasticity index that we calculated is predictive of the percentage change in leaf area under drought relative to well-watered conditions.

**Fig. S7.** The identification of our candidate genes upon application of several different GWAS methods improves confidence in candidates for leaf area plasticity and intrinsic water use efficiency (WUE_i_).

**Fig. S8.** Genotypic and transcriptomic reaction norms based on the strength of association of individual SNPs and transcripts highlight the variation in the strength of these associations depending on water availability.

**Fig. S9.** The genetic and transcriptomic basis of the relationship between leaf area plasticity potential in response to drought with (Figs S9A, B) leaf area potential and (Figs S9C, D) intrinsic water use efficiency (WUE_i_).

**Fig. S10.** Illustration of candidate transcript variants that regulate intrinsic water use efficiency in Iberian Arabidopsis accessions.

**Table P1.** Leaf area measurements recorded in this experiment (well-watered and drought). Calculated relative leaf area (rLA) and plasticity, water use efficiency (WUE_i_, δ^13^C) values recorded by Dittberner *et al*. (2018) and MOD17A2 (net primary productivity) information obtained from AraCLIM (Ferrero-Serrano & Assmann, 2019) for the accessions included in our study.

**Table P2.** Water use efficiency (WUE_i_, δ^13^C) values recorded by Dittberner *et al*. (2018) for 95 Iberian Arabidopsis accessions.

**Table G1.** Candidates obtained from GWAS on relative leaf area (rLA) under well-watered conditions. Only associations with a score ≥ 4 are included here. For a raw, unfiltered, as well as for a fully filtered and annotated (see Materials & Methods) version of this GWAS output, please visit the data repository.

**Table G2.** Candidates obtained from GWAS on relative leaf area (rLA) under drought. Only associations with a score ≥ 4 are included here. For a raw, unfiltered, as well as for a fully filtered and annotated (see Materials & Methods) version of this GWAS output, please visit the data repository.

**Table G3.** Candidates obtained from GWAS on leaf area plasticity. Only associations with a score ≥ 4 are included here. For a raw, unfiltered, as well as for a fully filtered and annotated (see Materials & Methods) version of this GWAS output, please visit the data repository.

**Table G4.** Candidates obtained from GWAS on water use efficiency (WUE_i_, δ13C) values recorded by Dittberner *et al*. (2018) for the accessions included in our study. Only associations with a score ≥ 4 are included here. For a raw, unfiltered, as well as for a fully filtered and annotated (see Materials & Methods) version of this GWAS output, please visit the data repository.

**Table G5**. Candidates obtained from GWAS on water use efficiency (WUE_i_, δ13C) values recorded by Dittberner *et al*. (2018) for 95 Iberian Arabidopsis accessions. Only associations with a score ≥ 4 are included here. For a raw, unfiltered, as well as for a fully filtered and annotated (see Materials & Methods) version of this GWAS output, please visit the data repository.

**Table G6.** Candidates obtained from eGWAS on the natural variation in transcript abundance of *SAUR26* extracted from Kawakatsu *et al*. (2016). For a raw, unfiltered, as well as for a fully filtered and annotated (see Materials & Methods) version of this GWAS output, please visit the data repository.

**Table T1**. Results from TWAS (r_s_).

**Table T2.** Results from TWAS (*P*-value).

**Table GO1.** GO analysis on the 31 genes corresponding to the top 50 candidate SNPs resulting from GWAS analysis on leaf area plasticity index for drought.

**Table GO2.** GO analysis on the 58 genes identified from TWAS to exhibit transcript abundances that are negatively correlated with leaf area plasticity index for drought, and negatively correlated with water use efficiency determined from (δ^13^C) values from accessions overlapping with our sample (red datapoints in Fig. 5C).

**Table GO3.** GO analysis on the 45 genes identified from TWAS to exhibit transcript abundances that are positively correlated with water use efficiency determined from (δ^13^C) values in 88 Iberian Arabidopsis accessions (datapoints with r_s_ ≥ 0.4 in Fig. 6B).

**Table GO4.** GO analysis on the 494 genes identified from TWAS to exhibit transcript abundances that are positively correlated with leaf area plasticity index for drought (green and grey datapoints with r_s_ ≥ 0.4 in the y-axis representing leaf area plasticity index in Fig. 5C).

**Table GO5.** GO analysis on the 28 genes identified from TWAS to exhibit transcript abundances that are negatively correlated with relative leaf area (rLA) under well-watered conditions, and positively correlated with leaf area plasticity index for drought (blue datapoints in S9B).

**Table GO6.** GO analysis on the 11 genes identified from TWAS to exhibit transcript abundances that are negatively correlated with water use efficiency determined from (δ^13^C) values in 88 Iberian Arabidopsis accessions (datapoints with r_s_ ≤ −0.4 in Figure 6B).

**Table GO7.** GO analysis on the 302 genes identified from TWAS to exhibit transcript abundances that are positively correlated with relative leaf area (rLA) under drought.

**Table GO8.** GO analysis on the 134 genes identified from TWAS to exhibit transcript abundances that are negatively correlated with relative leaf area (rLA) under well-watered conditions (blue and grey datapoints with r_s_ ≤ 0.4 in the x-axis of Figure S9B).

**Table GO9**. GO analysis on the 177 genes identified from TWAS to exhibit transcript abundances that are positively correlated with relative leaf area (rLA) under well-watered conditions (yellow and grey datapoints with r_s_ ≥ 0.4 in the x-axis of Figures S9B).

**Table GO10.** GO analysis on the 233 genes identified from TWAS that exhibit transcript abundances which are negatively correlated with relative leaf area (rLA) under drought.

## Acknowledgements

This research was supported by funding from The Pennsylvania State University to S.M.A. We thank Mr. Leland Burghard, Ms. Sarah Cronk, and Dr. Yotam Zait for greenhouse assistance. We thank Drs. David Chakravorty and Yotam Zait for helpful comments on the manuscript.

## Author contributions

S.M.A & A.F-S. conceived the project, research plans, and experimental design. A.F-S. performed the experiments while S.M.A provided guidance throughout the experimental work. A.F-S. analyzed the data and prepared the figures; S.M.A supervised data analysis and figure preparation. S.M.A & A.F-S wrote the manuscript.

## Data Availability

The data supporting the results are available as Supporting Information files. The code and data to easily reproduce the analysis, figures, and tables in this manuscript are available at https://github.com/AssmannLab and will be uploaded to Zenodo upon acceptance.

