## Supplementary material for "Genetic variation associated with plastic and homeostatic growth responses to drought in Arabidopsis": SI Figures

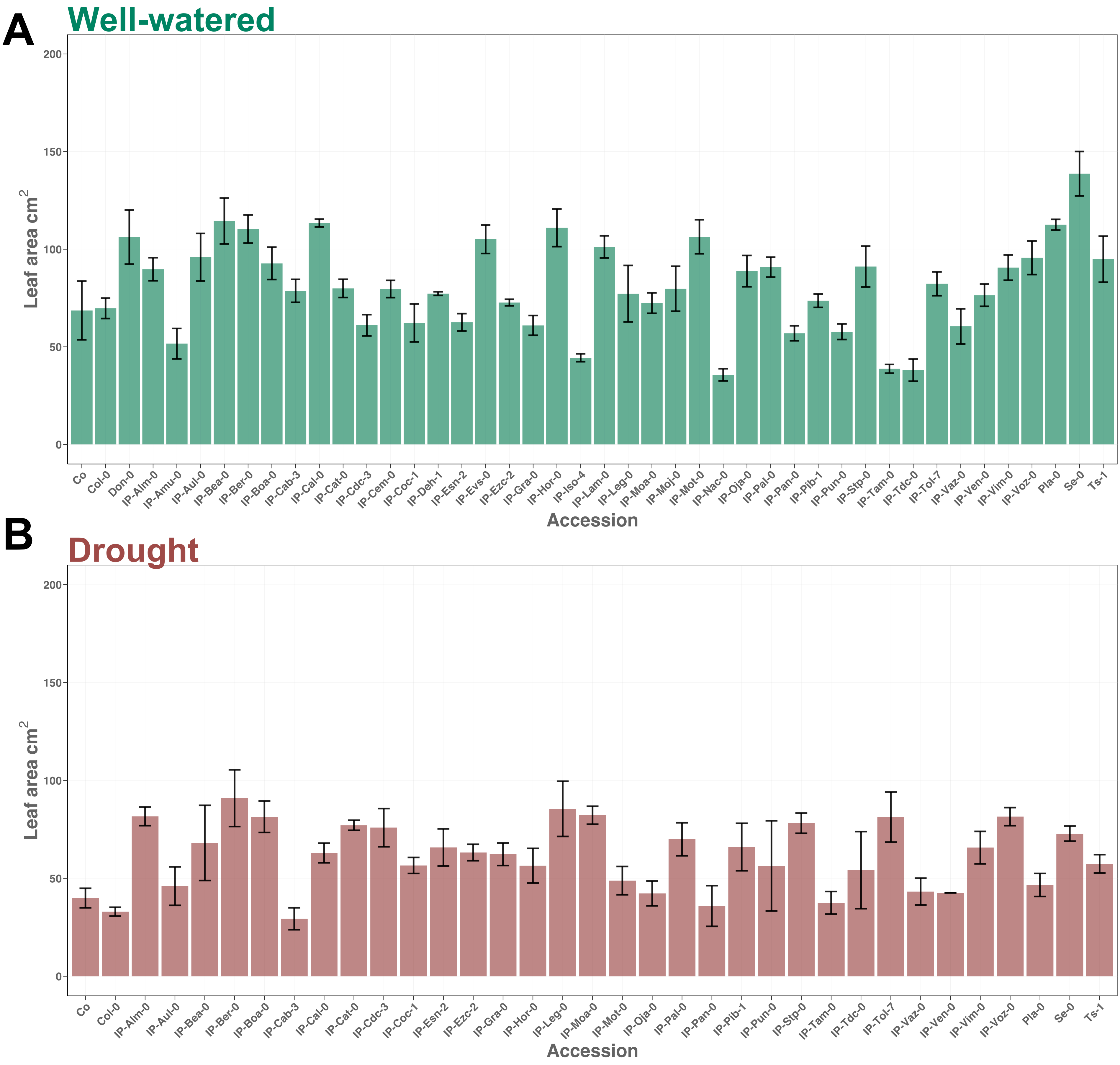


**Fig. S1.** Rosette leaf area observations at the time of flowering (Table P1).

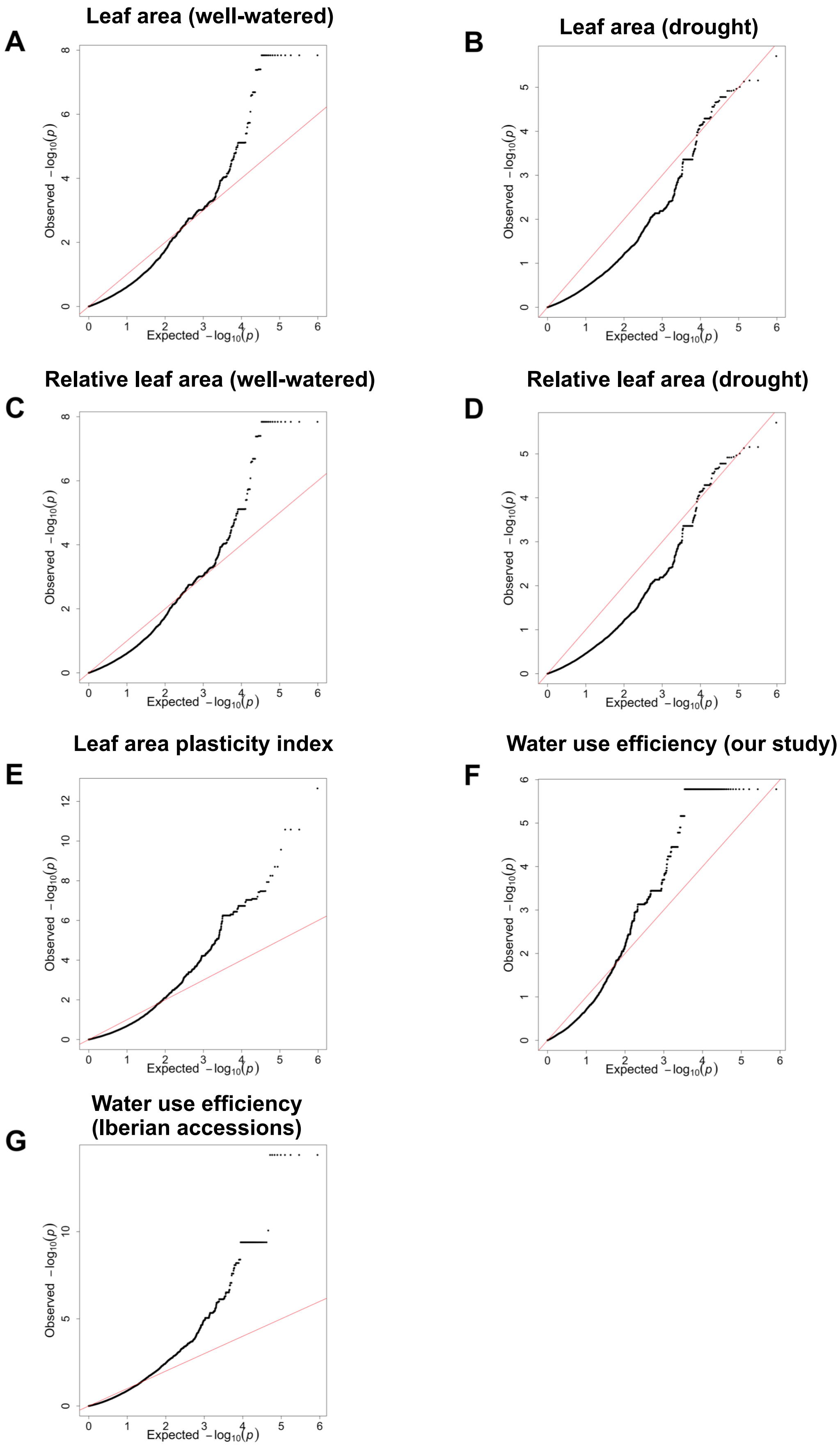


**Fig. S2.** Quantile-quantile (QQ) plots for GWAS analysis using the Linear Model.

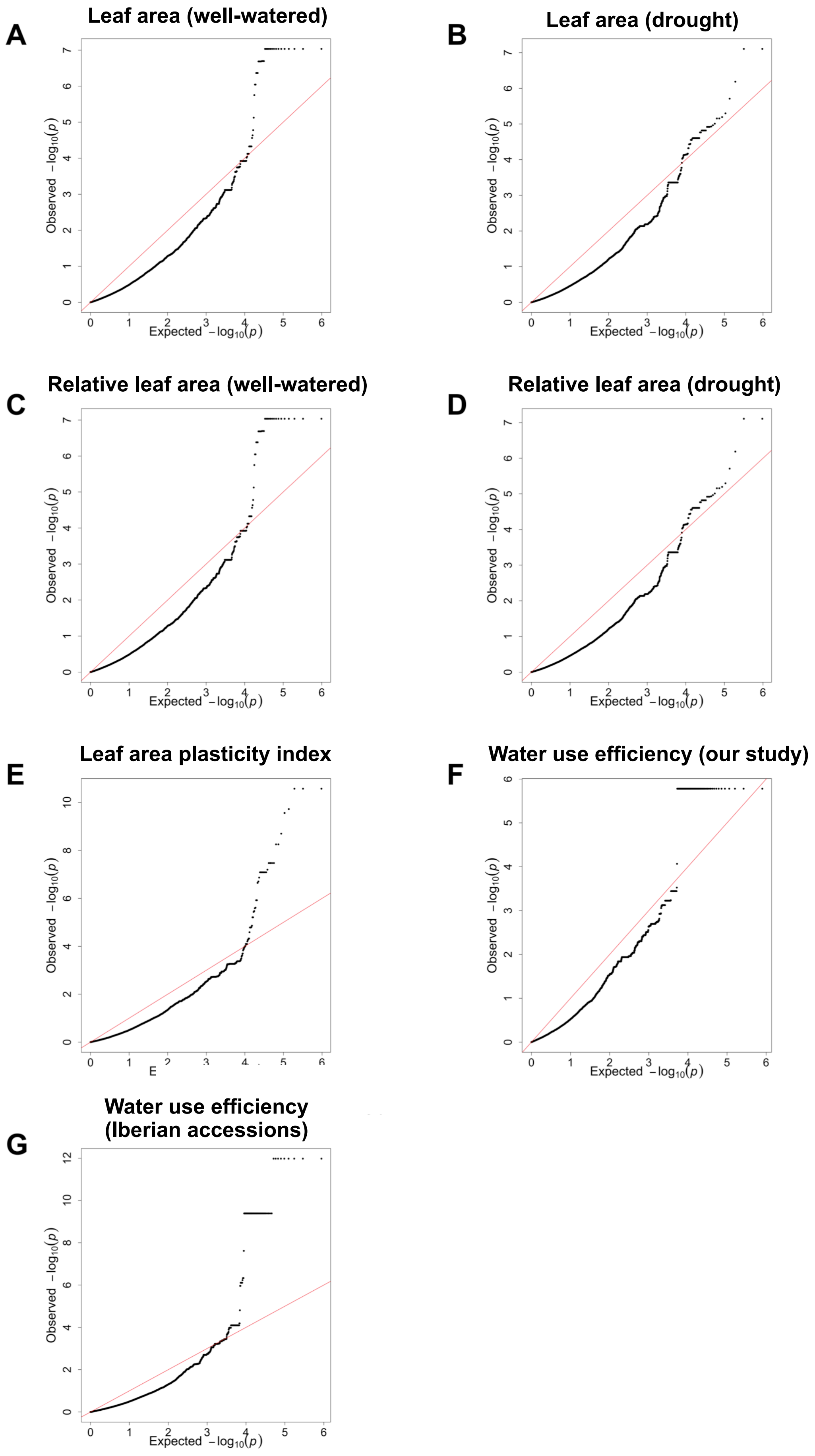


**Fig. S3.** Quantile-quantile (QQ) plots for GWAS analysis using the Accelerated Mixed Model.

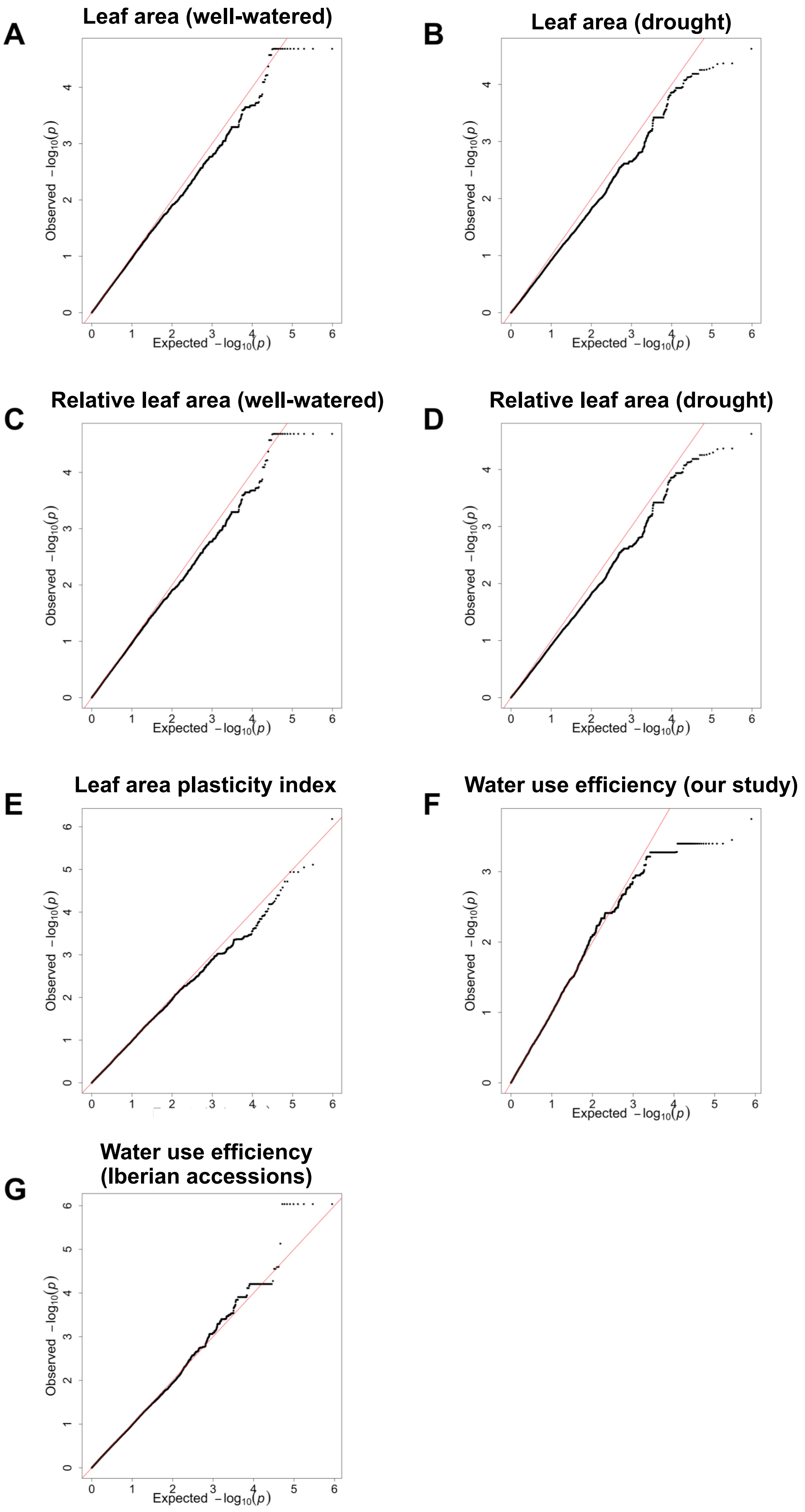


**Fig. S4.** Quantile-quantile (QQ) plots for GWAS analysis using GWAS-Flow.

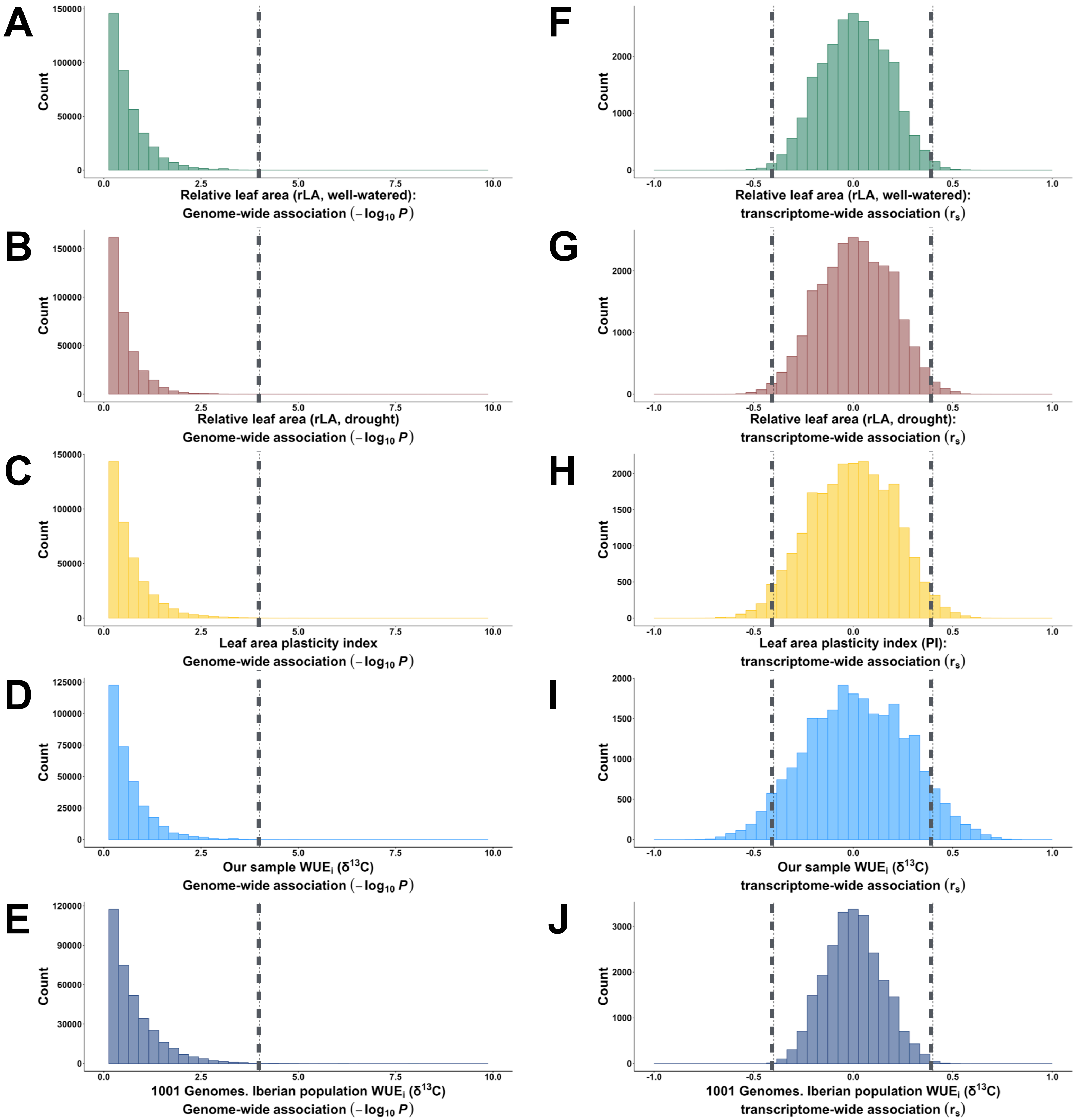


**Fig. S5.** Histograms of the distribution of $r_s$ and scores obtained from GWA and TWA analyses of the traits discussed in this study illustrate the stringent significance thresholds imposed in this study (vertical dashed lines).

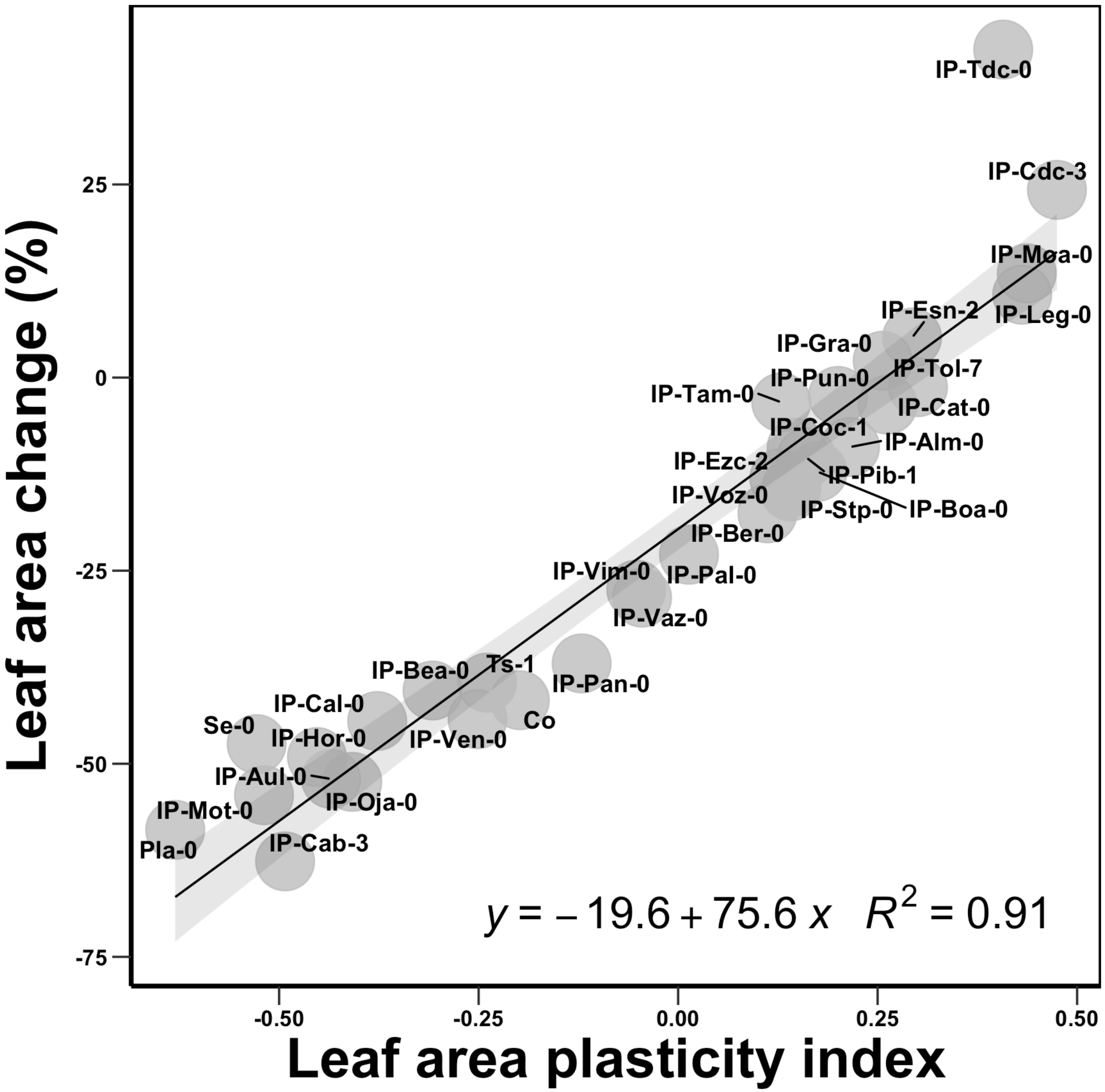


**Fig. S6.** The plasticity index that we calculated is predictive of the percentage change in leaf area under drought relative to well-watered conditions.

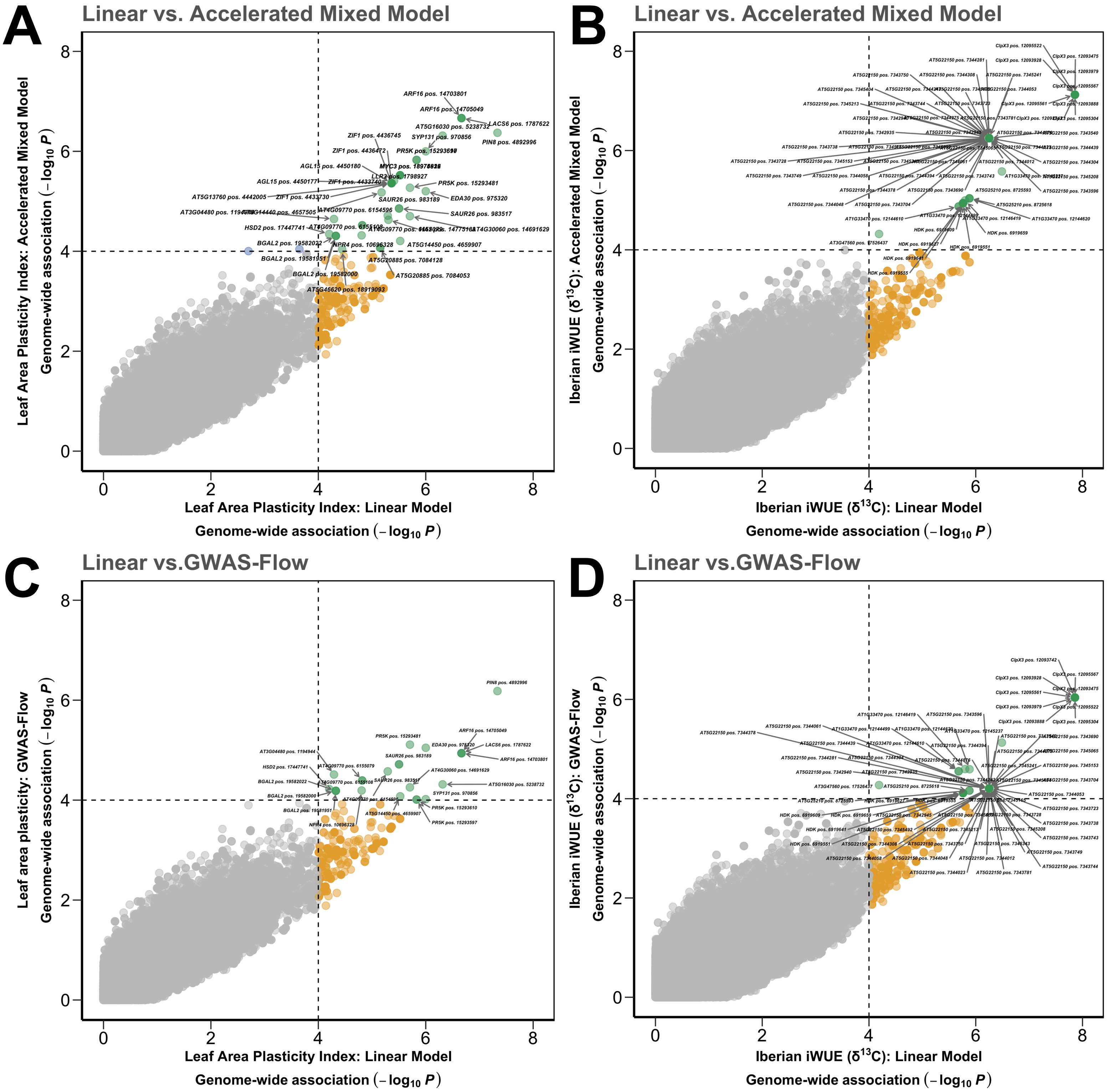


**Fig. S7.** The identification of our candidate genes upon application of several different GWAS methods improves confidence in candidates for leaf area plasticity and intrinsic water use efficiency ($WUE_i$).

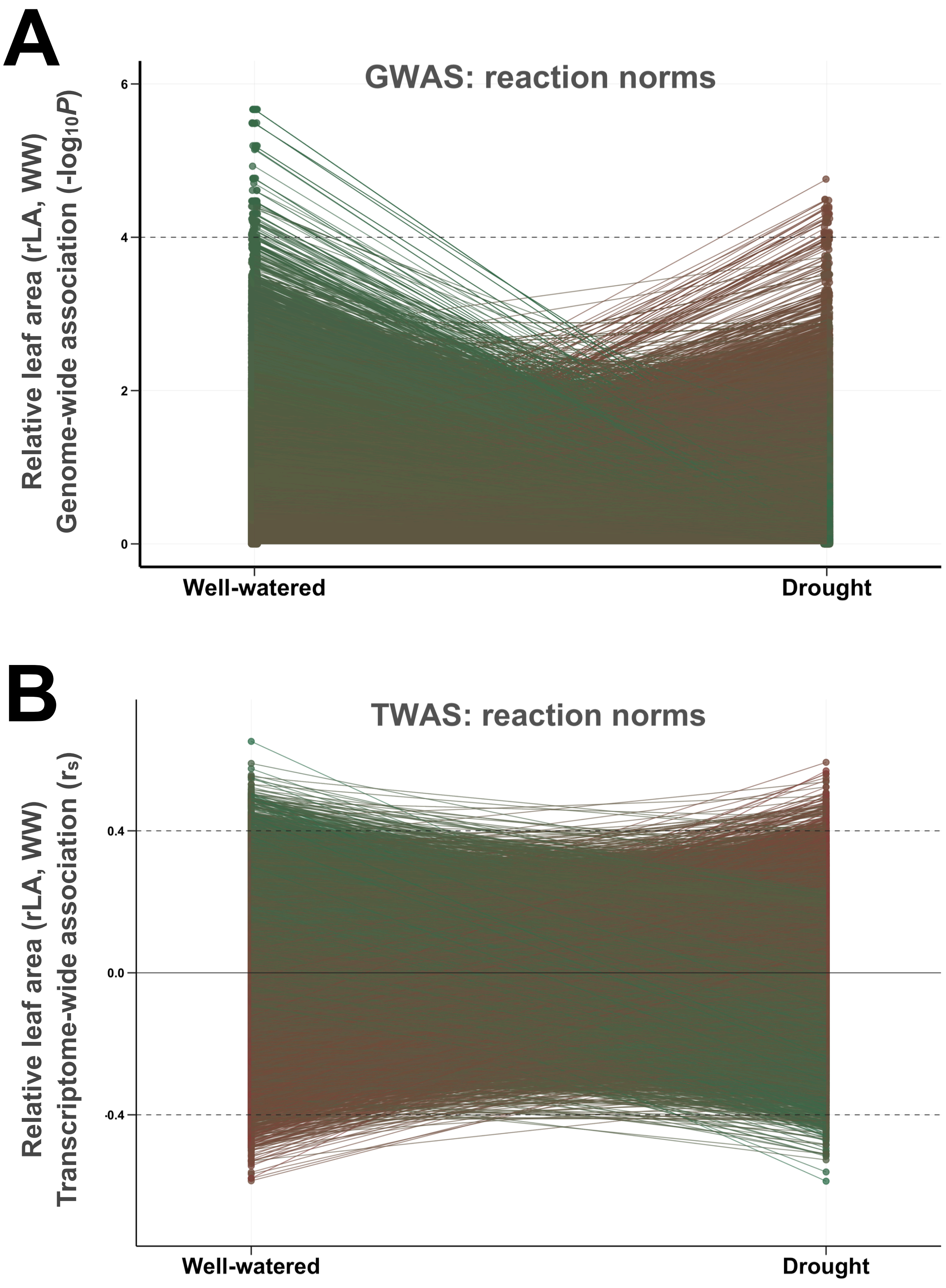


**Fig. S8.** Genotypic and transcriptomic reaction norms based on the strength of association of individual SNPs and transcripts highlight the variation in the strength of these associations depending on water availability.

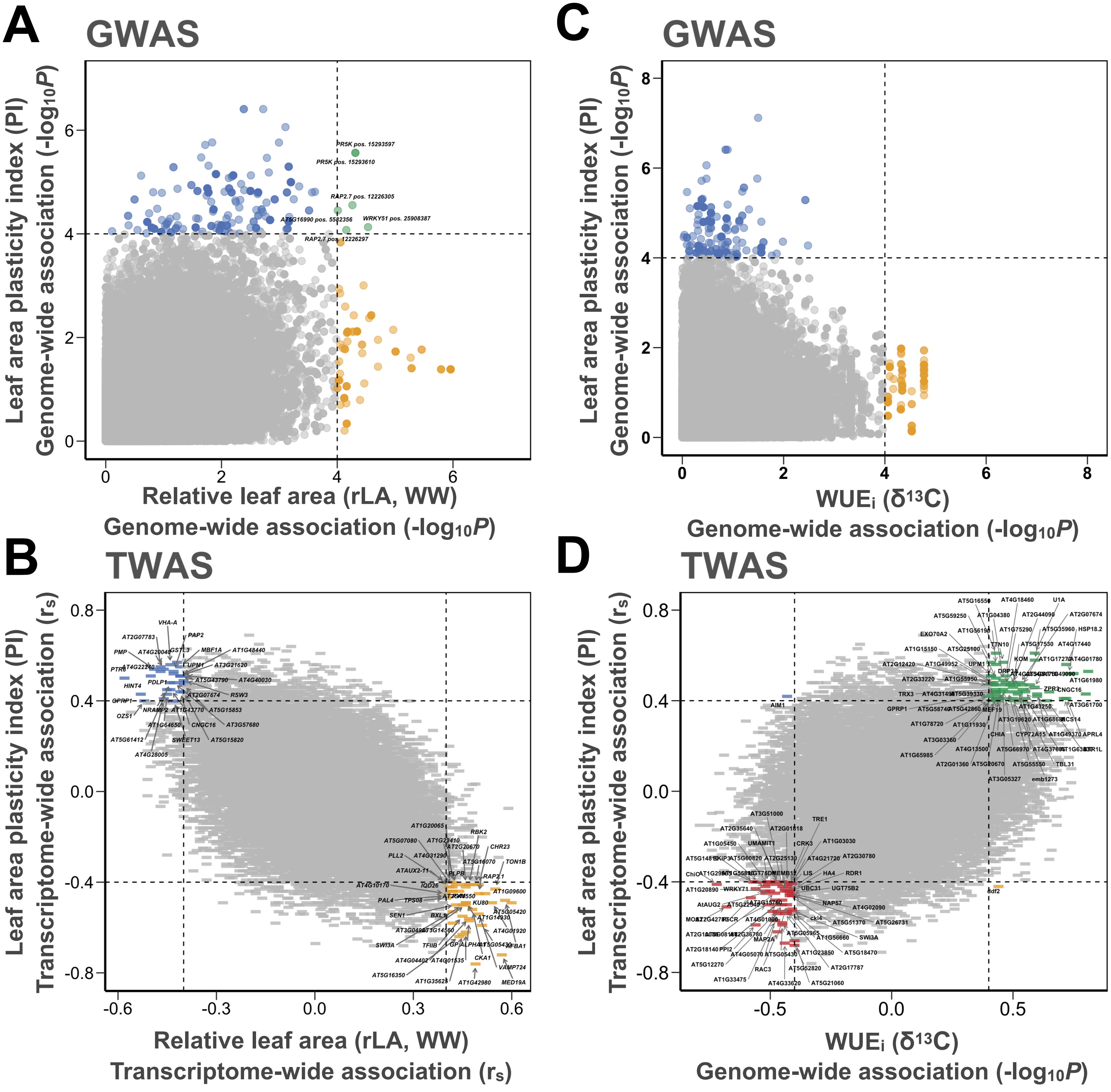


**Fig. S9.** The genetic and transcriptomic basis of the relationship between leaf area plasticity potential in response to drought with (Figs S9A, B) leaf area potential and (Figs S9C, D) intrinsic water use efficiency (WUEi).

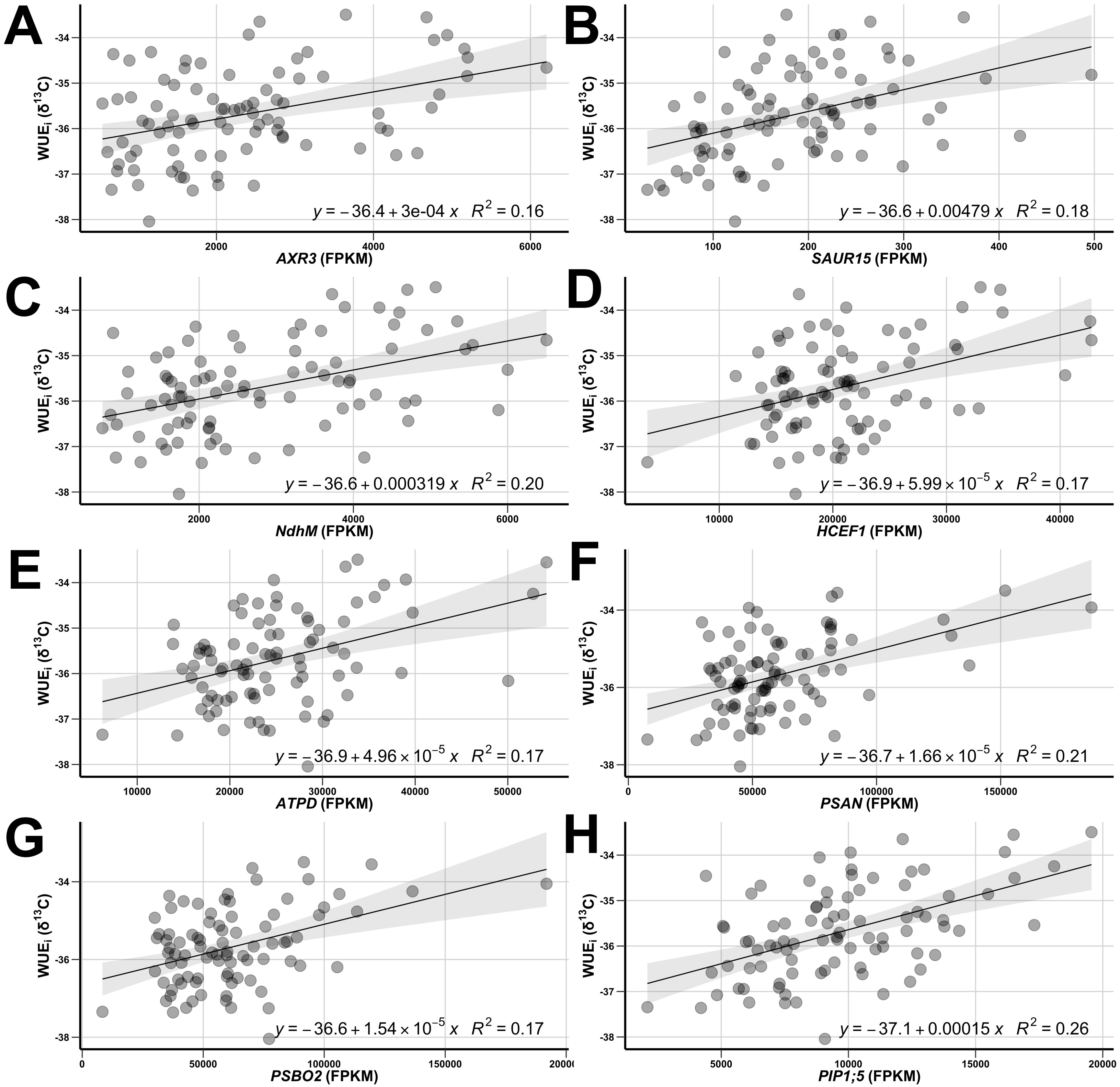


**Fig. S10.** Illustration of candidate transcript variants that regulate intrinsic water use efficiency in Iberian Arabidopsis accessions.
