## Supplementary material for "Genetic variation associated with plastic and homeostatic growth responses to drought in Arabidopsis": Document S1

### Challenges, caveats, and limitations

Determining the genetic basis of drought adaptation in studies of natural variation is not trivial. The distribution of complex phenotypic traits and their plasticity in a population is often continuous (Galton, 1889) because they are influenced by variation in a large number of genes. The effect of many of these variants is additive or synergistic and of small effect individually (Fisher, 1919), and thus difficult to identify (Barton *et al.*, 2019; Sohail *et al.*, 2019). For example, leaf area displays natural intraspecific variation and varies in a continuous manner in response to resource availability. Leaf area is an important trait in relation to carbon and water fluxes: the degree of growth and leaf area production indicates the total amount of light intercepted by plants, assimilated carbon, but also water availability and evaporative water loss.

Our study has revealed genetic variants associated with both plasticity and homeostasis in leaf area following drought. One limitation of our study is that while the number of accessions that we utilized would have been considered ample just a few years ago, with the more recent advent of GWA studies in Arabidopsis such studies typically include more accessions. Given the reduced power X If the number of accessions utilized is low, low-frequency variants may not be present in the subset of accessions studied, and thus the full genomic signature of a trait may not be captured. This is less of a concern when we focus on variants of large effect that are undergoing selection and thus are at higher frequency in the population. Accordingly, in this study, we imposed a stringent threshold and did not consider variants in frequencies lower than 10% as we were interested in adaptive genetic variation. In addition, we performed TWAS, which is less susceptible to this concern because it evaluates continuous variation in transcript abundance rather than the presence/absence of SNPs within the study population. Indeed, RNA-seq experiments often are conducted on very few genotypes, frequently comparing transcript abundance between just two genotypes: wild-type and a specific mutant of interest. Another limitation typical of GWAS is the fact that synthetic associations, non-causative markers that are within linkage disequilibrium with causative makers, are difficult to discern from true causative markers (Korte and Farlow, 2013; Sasaki et al.*et al.*, 2021). Identification of polygenic association signals through GWAS is also sensitive to bias introduced by uncorrected population structure that can also introduce a number of false positives in the resulting list of candidates (Barton et al.*et al.*, 2019; Sohail *et al.*, 2019). There are statistical methods to address these issues, but such methods also introduce a number of false negatives due to over-correction (Korte and Farlow, 2013). The need to correct for confounding effects of population stratification, family structure, and cryptic relatedness can be minimized by focusing on regional collections, such as the Iberian population in this case (Frachon *et al.*, 2018; Tabas-Madrid *et al.*, 2018). Moreover, while we illustrate our analysis using a linear model that does not correct for population structure, we include the same analyses but corrected for population structure through use of an Accelerated Mixed Model approach using both GWAPP (Seren et al.*et al.*, 2012) and GWAS-Flow (Freudenthal *et al.*, 2018) (Tables G1-G6). While we discuss our results using the outputs from the linear model, it is important to point out that the individual candidate variants discussed in this study are significant for all three analyses, which greatly improve confidence in them (Tables G1-G6).

TWAS analysis does not suffer from the limitations imposed by false non-causal synthetic associations due to linkage disequilibrium (Li *et al.*, 2021a) typical of GWAS (Korte and Farlow 2013). However, TWAS analysis also presents a series of challenges, as this method is sensitive to time and tissue-dependent expression (Li *et al.* 2020b; Wainberg *et al.* 2019). The combined analysis of reference genomic (The 1001 Genomes Consortium 2016) and transcriptomic data (Kawakatsu *et al.* 2016), as done here, can partially ameliorate the limitations of each method.

The validation of candidates from both GWA and TWA studies also suffers from the challenge of restricted genetic backgrounds. Most often, the characterization of knock-out mutants of candidate genes is performed in a single genetic background, typically Col-0. Here, we instead validated our results by meta-analysis of the larger Iberian population. TWAS in this larger sample identified 31 out of the 45 candidate genes that we previously identified in the subset of 20 Iberian accessions in our study for which published $\delta^{13}C$ data were available (Table T1). Moreover, there are significant correlations between the results from our sample and the larger population from both GWAS and TWAS (Fig. S8), indicating that our sample choice is representative of the larger population.

For the associations between the phenotypic variables we obtained in this study, and the local environment, we inspected the data using Cook's distance (Cook, 1984; Altman and Krzywinski, 2016) to evaluate the influence of each datapoint on the fit. We considered values greater than 4/n to be overly influential. Based on this, we decided to remove the phenotypic values we obtained for the accession IP-Coc1 (accession id = 9535), which were consistently unrealistically influential across different phenotypes based on their extreme Cook's distance.

We include these removed values in Table S1. It is true that perhaps these overly influential could have facilitate the identification of associated variants of large effect that we may not obtain when this outlier is removed. On the other hand, this outlier may just reflect unrealistic values that could have affect the results. We opted for a conservative strategy and removed these outliers

for genome-wide association mapping in Arabidopsis. *Plant Cell* **24:** 4793–4805.

**The 1001 Genomes Consortium** 2016. 1,135 Genomes reveal the global pattern of polymorphism in *Arabidopsis thaliana*. *Cell* **166:** 481–491.

**Wainberg, Michael, Nasa Sinnott-Armstrong, Nicholas Mancuso, Alvaro N. Barbeira, David A. Knowles, David Golan, Raili Ermel *et al.* 2019.** Opportunities and challenges for transcriptome-wide association studies. *Nature genetics*. **51:** 592-599.
